# Targeting plasma membrane and mitochondrial instability in breast cancer cells and breast epithelial to mesenchymal transition-model cells by adamantyl diaza-crown ether ZG613

**DOI:** 10.1101/2022.09.30.510273

**Authors:** Katja Ester, Marija Mioč, Pavel Spurny, Daniel Bonhenry, Marko Marjanović, Lidija Uzelac, Jelena Gabrilo, Tatjana Šumanovac, Wolfgang Schreibmayer, Kata Majerski, Babak Minofar, Jost Ludwig, Marijeta Kralj

## Abstract

The adamantane derived diaza-crown ether ZG613 was assessed as a potential breast cancer cells and breast epithelial to mesenchymal transition (EMT)-model cells targeting agent. We postulated that ZG613 activity relies on its plasma/mitochondria membrane disruption ability based on adamantane hydrophobicity and/or crown ether related ionophoric properties. We performed molecular dynamics (MD) simulations and next generation sequencing, followed by *in vitro* study of cell death, membrane perturbations and ionophoric ability, as well as *in vivo* study of effects on the tumour growth. MD simulation and RNA sequencing pointed toward physical disruption of plasma membrane by ZG613, corroborated by measured increase in membrane permeability leading to cell death. Measurements of ion fluxes confirmed ZG613 inability to transport Na^+^ and K^+^, as predicted by MD simulation. EMT-model cells exhibit changes in mitochondrial morphology and ATP levels, successfully targeted by ZG613. ZG613 caused mild retardation of tumour growth in vivo. In conclusion, ZG613 kills breast cancer cells and breast EMT-model cells by physical disruption of plasma membrane and impairments of mitochondrial functions. Breast EMT cells represent good potential targets within the breast tumour, due to their plasma membrane and mitochondrial instability.

## 1. Introduction

Maintenance of membrane integrity is vital for eukaryotic cell survival. Rapidly upon membrane breach, which can be induced by numerous factors including synthetic manipulators of membrane structure and composition, cells employ adopted protective mechanisms to avoid death^1,2^. Membranes in cancer cells have changed structure and function compared to healthy cells of the same lineage. Alterations include disturbances in plasma/organelle membrane’s lipid profile and fluidity^3–5^ as well as disturbances in bioelectric properties based on changes in both plasma^6,7^ and mitochondrial membrane potential^8,9^. Cancer membrane targeting is currently developing as an effective therapeutic strategy^10–12^. Antimicrobial drugs with ionophoric properties as salinomycin^13–15^, monensi^16,17^, azurin^4^ and gramicidin ^18,19^, as well as amphiphilic drugs with potential to locally perturb composition of lipid bilayers throughout the cell^11,20^, have been repurposed to target membranes in cancer. One of the most promising approaches involves drugs with heterogeneous anticancer activity that simultaneously hit multiple membrane-related targets within the cell. Multiple targets include contributors of Multidrug resistance (MDR), ion channels, membrane receptors and lipidomic homeostasis^10,21^.

Breast cancer is the second most common cause of death from cancer in woman. This heterogeneous disease is comprised of different types, the most hazardous being characterised by poor prognosis and inefficient chemotherapy. Breast cancer chemoresistance is often related to a high percentage of cancer stem cells (CSCs) within the tumour bulk, especially in triple negative breast cancer (TNBC) CSCs, also named tumour-initiating cells, share some characteristics with healthy stem cells, such as pluripotency and self-renewal, while they can survive chemo and radiotherapy and give rise to a new tumour^23^. CSCs subpopulation within a tumour overlaps with a subpopulation of cells that have undergone epithelial to mesenchymal transition (EMT), developmental program activated during cancer invasion and metastasis^24–26^. EMT is characterised by remodelling of cytoskeleton, changes in plasma membrane composition and modifications of cellular organelles, metabolic reprogramming and other adopted traits that increase survival of invasive cell^27–29^. Regarding membrane biophysics, EMT-related changes cause enhancement in plasma membrane fluidity^30–32^. In addition, important steps for induction and maintenance of EMT include interplay between cytoskeleton and mitochondrial trafficking^33^ and induction of mitochondrial fusion^34^. Due to the growing need to study breast CSCs, cell lines that are induced to undergo EMT and therefore acquire CSC-like phenotype were designed^24^. EMT-model cells enable screening of drugs that target chemoresistant subpopulations among tumour bulk. Promising agents delivered by this approach include salinomycin^13^ and combinatorial therapy for TNBC, the mesenchymal stem cancer^25^.

Crown ethers are heterocyclic compounds containing cyclic polyethers. Although crown ethers are widely used, their biomedical applications rely mostly on their potential for ion transport^35–37^. In our previous research, we performed the first systematic study of the antitumor effects of crown and aza-crown derivatives^38^, where we demonstrated potent antitumor activity of crown ethers. In the following study, computational structure activity relationship analysis of oxa-, monoaza-, and diaza-18-crown-6 derivatives revealed that hydrophobicity of the attached substituents and side-chains represents the main determinant of bioactivity^39^. Based on ALOGP non-linear regression model prediction, we synthesised six novel adamantane-containing crown ethers. Compound ZG613, adamantane-functionalized diaza-bibracchial lariat ether (Fig. 1) has shown the highest biological activity, and additionally, exhibited best cation extraction ability^40^, prompting us to choose it for further analysis. In a parallel study, we tested the potential of adamantane-substituted aza and diaza-18-crown-6 derivatives to inhibit a major MDR contributor, P-glycoprotein (MDR1)^41^, where ZG613 indirectly inhibited P-glycoprotein-mediated efflux of drugs, possibly by physical interference with plasma membrane. Based on our published data we postulated that ZG613 cytotoxicity relies on the adamantane hydrophobicity. In addition, the compound acts as potassium/sodium ionophore via formation of stable complexes with alkali metal cations. Thus, we investigated the potential of adamantyl crown ether to disturb both lipid and ion homeostasis within the tumour cell. An important strategy to target cancer cells is to destabilise plasma/organelle membrane (already impaired in tumours and EMT). Hence, we reasoned that ZG613, the compound with membrane impairment potential, would successfully target breast cancer cells and breast EMT-model cells.

**Figure 1.**
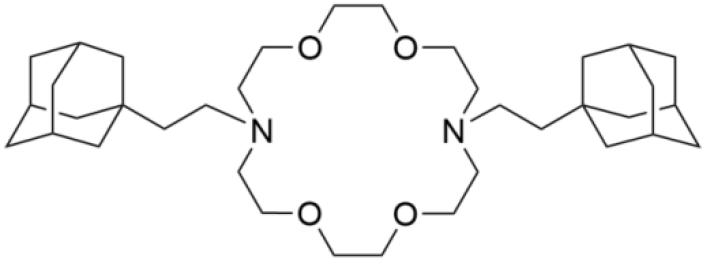
*N,N*’-bis[2-(1-adamantyl)ethyl]-4,13-diaza-18-crown-6

## 2. Material and Methods

### 2.1. Molecular dynamics simulations

Parameters for the crown ethers were derived by using antechamber^42^. The general amber force field (GAFF)^43^ has been used to create the topology of these molecules. The TIP3P water model^44^ was used to describe water molecules. Charges and Van der Waals parameters for the ions are derived by using a rescaling scheme and a mean field approach^45^. Amber parameters were used for the chloroform molecules. Finally, the Slipids force field^46^ was used for the phospholipids. Configurations for the membrane and equilibration steps were obtained from the CHARMM-GUI web-server^47,48^. Crown ethers were added to the system by putting them randomly in bulk water using PACKMOL^49^. The input files were then updated for the SLipids force field. In total, eight ZG613 molecules were added to the membrane which is constituted by two leaflets of 147 phosphatidylcholine (POPC) molecules, each hydrated by roughly 19000 water molecules with an ionic strength corresponding to 0.15 M KCl. The minimization and equilibration steps were kept^50^. Production was then run for 100 ns. Molecular dynamics simulations were all performed with GROMACS^51^ (version 2018). A time step of 2 fs was used for all simulations. A leap-frog algorithm for integrating Newton’s equations of motion was used. The LINCS^52^ algorithm was used to constraint all covalent bonds. For membrane simulations, the NPT ensemble was chosen. The pressure was kept constant at 1.013 bar by using the Parrinello–Rahman barostat^53^ with a coupling constant of 10.0 ps and an isothermal compressibility of 4.5 × 10^-5^ bar^-1^. A semi-isotropic scheme was used where the pressure in the xy directions is decoupled from the direction normal to the bilayer (z direction). The Nosé–Hoover thermostat^54,55^ was used to keep the temperature constant at 300K with a coupling constant of 0.5 ps. The water with the ions, membrane and crown ethers were coupled to separate thermostats. Long-range electrostatics were treated with the particle mesh Ewald scheme^56,57^. The real-space cut-off was 1.4 nm with a Fourier spacing of 0.12 nm. The cut-off for Lennard-Jones interactions was set to 1.4 nm and the isotropic long-range corrections were applied to energy and pressure. PACKMOL was used to create the initial configurations for the water/chloroform systems. To create the biphasic system, 2000 water molecules were used to create a slab of water while another slab containing 750 molecules of chloroform was added. The initial dimension of the box was 4 nm × 4 nm × 8 nm. The energy minimization was then performed on these initial configurations, using the “steepest descent” method. A canonical ensemble (NVT) was employed for 100 ps to equilibrate the system at the required temperature using a V-rescale thermostat^58^ with a time constant of 0.1 ps, following NPT simulations on the output of the NVT step for 20 ns. The pressure was controlled by the Parrinello-Rahman barostat. X and Y components were decoupled from Z (the component normal to the interface) using a semi-isotropic coupling. Equilibration was monitored by calculating the density of chloroform (1.48 g/cm3 at 298.15K) and water. All simulations are visualized using the visual molecular dynamics (VMD) software (version 1.9.3)^59^.

### 2.2. Cells and reagents

The synthesis of adamantane derived diaza-crown ether ZG613 was previously described^39^. Prior to each assay, the compound was freshly dissolved in DMSO using 2 min sonication in an ultrasonic bath.

HMLE cells transduced with lenti-Twist (HMLE-Twist) and HMLE cells transduced with an empty control vector (HMLEpBp), were kindly provided by Dr. Robert A. Weinberg and Tamer T. Onder. HMLE cells were maintained in a 1:1 mixture of HuMEC Ready Medium (Gibco, NY, USA) and DMEM complemented with 10% FBS, 2 mM L-glutamine, 100 U/mL penicillin, 100 μg/mL streptomycin, 2.5 μg/ml insulin and 0.5 μg/mL hydrocortisone (Sigma-Aldrich, MO, USA). SUM 159 triple-negative breast cancer cells were kindly provided by Dr. Robert A. Weinberg and were maintained in Ham’s F-12 media with L-Glutamine and Pen/Strep, with 5% FBS, 5 μg/ml Insulin (Sigma-Aldrich) and 1 μg/ml Hydrocortisone. MCF7 breast adenocarcinoma cell line was obtained from ATCC and maintained in DMEM complemented with 10% FBS, 2 mM L-glutamine, 100 U/mL penicillin and 100 μg/mL streptomycin. MJ90 (HCA2) fibroblasts, isolated previously from neonatal foreskin, were kindly provided by Dr. I. Rubelj and were maintained as MCF7. All cells were grown in a humidified atmosphere at 37 °C with 5% CO_2_. EMT and non-EMT status was regularly checked by probing for EMT markers (CD24/CD44 by FACS and E- and N-cadherin by western blot).

Digitonin, FCCP, and DiBAC were purchased from Sigma-Aldrich. Flow-cytometry reagents were from BD Bio sciences (San Jose, California, USA). All of the remaining reagents, if not stated otherwise, were purchased from Sigma (Sigma Aldrich, MO, USA) or Kemika (Zagreb, Croatia).

### 2.3. Antiproliferative activity and cell death measurements

#### 2.3.1. Cell viability

Cell viability assay was performed as described previously according to the modified procedure of the National Cancer Institute, Developmental Therapeutics Program^60^. Cells were seeded at 1-5 × 10^3^/well (depending on the doubling time of a specific cell line) in standard 96-well microtiter plates and left to attach for 24 hr. Next day, the compound was added in five serial 10-fold dilutions. The final concentration of DMSO was < 0.2% which was non-toxic to cells. The cell growth was evaluated after 72 hours of incubation, by adding MTT reagent (Sigma-Aldrich), which measures the reduction of a tetrazolium component (MTT) into an insoluble formazan product by the mitochondria of viable cells. After four hours of incubation the precipitates were dissolved in DMSO, and the absorbance (A) was measured on Multiskan EX (Thermo Scientific) Plate Reader at 570 nm. The percentage of growth (PG) of the cell lines was calculated according to one or the other of the following two expressions:

If (A_test_ –A_tzero_) ≥ 0 then:

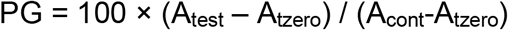

If (A_test_ – A_tzero_) < 0 then:

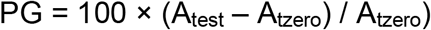

Where:

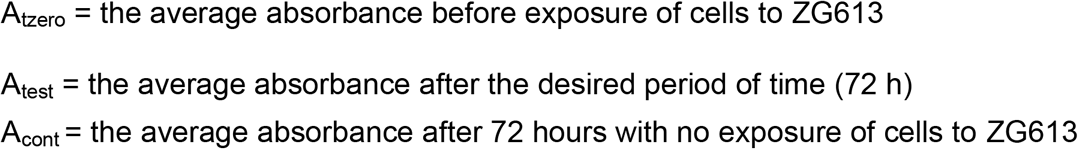

Each test was performed in quadruplicate. The results from at least three individual experiments are expressed as dose-response curves (Figure 3A). A negative percentage indicates cytotoxicity following drug treatment, where −100% shows no cells survived the treatment at the specific drug concentration. The results are also expressed as IC_50_, the concentration necessary for 50% of inhibition. The IC_50_ values were estimated in Excel from dose-response curves using linear regression analysis by fitting the test concentrations that give PG values above and below 50.

#### 2.3.2. Modality of cell death

To elucidate modality of cell death, modified MTT assay was performed. Cells were seeded at 1-5 × 10^3^/well in 96 well plates. The next day, cells were treated for 24 hr with an inhibitor of apoptosis Z-VAD-FMK (ALX 260020, Enzo, NY, USA) or inhibitor of necroptosis Necrostatin-1 (BML-AP309, Enzo, NY, USA). On the third day, cells were incubated for additional 24 hr with 5 μM ZG613 or vehicle (DMSO). Cells were treated with ZG613 alone or in the combination with inhibitors, as indicated in Results. Percentage of growth was calculated as experimental values divided by control values obtained with cells only treated with the solvent (DMSO).

#### 2.3.3. Lactate Dehydrogenase (LDH) Release

Cells were seeded at 5 × 10^3^/well into 96 well plates. The next day, serial two-fold dilutions of ZG613 were added and cells were incubated for various periods of time as indicated in the Results. Release of LDH was assessed by LDH CytoTox 96® Non-Radioactive Cytotoxicity Assay (G 1780, Promega, WI, USA) according to the manufacturer’s protocol. LDH positive control from the kit was used for Maximum LDH Release measurements as recommended. Absorbance (OD 490) was measured using Multiskan EX (Thermo Scientific) Plate Reader. Relative LDH Release was calculated as Experimental LDH Release divided by Maximum LDH Release.

### 2.4. RNA-Sequencing

Cells were seeded onto 10 cm plates (1 × 10^6^/plate) and left to attach overnight. Next day, media was replaced by media containing vehicle (DMSO) or 1 μM ZG613. After 24 hr, total RNA from biological triplicates of HMLE-Twist and HMLE-pBp cells was extracted with Quick-RNA Miniprep kit (Zymo Research) according to manufacturer protocol. DNase I was used to remove possible DNA contamination. RNA concentration was measured on Qubit 3.0 Fluorometer (Thermo Fisher Scientific) and the quality of RNA was additionally checked with Bioanalyzer 2100 (lowest RIN = 9.7, Agilent). cDNA library was prepared with TrueSeq Stranded mRNA kit (Illumina) from 750 ng of RNA (average cDNA size was ~345bp). Library was sequenced on NextSeq500 (Illumina) using high output flow-cell in paired-end mode (75 + 75 cycles). Raw data yielded over 690 million high-quality reads (Q30 > 75%). Reads were uploaded to BaseSpace Sequence Hub (Illumina) and aligned to human reference genome (hg19) via RNA-Seq Alignment app (STAR alignment, ver 1.1.1). Differential expression of genes after treatment for each cell line was executed via DESeq2 app (ver 1.1.0) in paired mode. Venn diagrams were plotted via web tool of Spanish National Biotechnology Centre^61^.

### 2.5. Gene Ontology

For analysis of Gene Ontology, we used The PANTER Classification System^62^ by loading single ranked genes with fold change > 1.1 from differential gene expression experiments. Up-regulated and down-regulated genes were separately loaded. Significant enrichment was determined by PANTER Statistical overrepresentation test via the Fisher’s Exact test and FDR correction method. GO’s were ranked using fold Enrichment Client Text Box Input.

### 2.6. Membrane permeability analysis

Membrane permeability changes were assessed by measurements of fluorescent dye SYTOX® Green (S7020, Thermo Fisher Scientific, USA) influx. Cells were plated at 1 × 10^4^/well in black 96-well plates using Bio Whittaker HBSS (BE10-527F, Lonza, Switzerland) with 4.5 g/L glucose. After initial 15 min equilibration with dye, ZG613, DMSO or non-ionic detergent Triton X (0.3%) were added immediately before measurements. Fluorescence was measured at absorption/emission maxima 485 nm/530 nm every 10 minutes during 120 min using Tecan M200 microplate reader. To calculate Sytox influx, average fluorescence values in each time point were divided by maximal value obtained with Triton X.

### 2.7. Analysis of mitochondrial structure and function

#### 2.7.1. Mitochondrial membrane potential

Cells at 5 × 10^5^/well in 6-well plates were treated with ZG613 in media at indicated concentrations. After 30 minutes of incubation, 2.5 μg/mL of mitochondrial potential-sensitive fluorescent dye JC-1 (eBioscience, EBI-65-0851-38) was added to the cells for additional 30 minutes of incubation (1h total of ZG613 treatment). Cells were then placed on ice and analysed on a BD FACSCalibur flow cytometer, and percentages of red and green fluorescent specific populations were calculated in FlowJo (TreeStar, Inc.).

#### 2.7.2. Mitochondrial morphology/Confocal imaging

For mitochondrial staining, cells were seeded on coverslips inside a 24-well plate in the concentration of 4 × 10^4^/well and treated with 5 μM ZG613 for 24h. Cells were washed in PBS, and MitoTracker Deep Red FM dye (Thermo Fisher Scientific, USA) was added at the concentration of 200 nM in PBS. The samples were incubated for 15 min under growth conditions, after which the MitoTracker solution was removed and the cells were rinsed thoroughly with PBS. Cells were fixed in 4% paraformaldehyde for 10 min and permeabilized with cold acetone for 5 min to improve the signal-to-background ratio. The coverslips were washed tith PBS and mounted using mounting medium with DAPI (Fluoroshield, Sigma-Aldrich, USA). Confocal fluorescence microscopy was performed using Leica TCS SP8 X microscope equipped with a HC PL APO CS2 63 × /1.4 oil objective (Leica Microsystems, Germany), with excitation/emission wavelengths of 644 nm/665–727 nm.

#### 2.7.3. Depletion of cellular ATP

Cells were plated at 2 × 10^4^/well in white 96 well plates, left for 2 hr to attach, and treated with ZG613 or DMSO for 30 min. Relative levels of ATP were assessed using CellTiter-Glo® Luminescent Cell Viability Assay (67572, Promega, WI, USA) according to the manufacturer’s protocol. Luminescence was measured using Tecan M200 Plate Reader. Levels of ATP were normalised to DMSO-treated control values.

### 2.8. Analysis of ion transport

#### 2.8.1. Electrophysiological recording and analysis

Cells were seeded on round glass coverslips in 24 well plates two days before the electrophysiological experiments. Before recording, the coverslips were transferred to a bathing chamber in bathing solution (NaBS) mounted to the stage of an inverted microscope (IM35, Zeiss, Germany). We performed whole cell patch clamp method and nystatin (N6261, Merck, Darmstadt, Germany) perforation as previously described^63^. Briefly, after Gigaseal formation, perforation was monitored by capacitance measurements. After successful perforation, as judged by capacitance > 25 pF, resting membrane potential was low pass filtered at 50Hz and recorded for 20 seconds using Axopatch-200A amplifier (Molecular Devices, USA). Traces were digitized at 1 kHz using the Digidata 1322A interface (Molecular Devices, USA) using Clampex 9.2 software (Molecular Devices, USA). Membrane resting potential was measured by averaging the entire trace using the Clampfit 10.3 software (Molecular Devices, USA). The composition of solutions used was as follows: NaBS: 2 mM KCl, 148 mM NaCl), 4 mM MgCl_2_, 1 mM CaCl_2_ and 10 mM HEPES, buffered with NaOH to pH:7.4. PFS: 20 mM KCl, 120 mM potassium-aspartate, 10 mM NaCl, 4 mM MgCl_2_, 1 mM EGTA-potassium salt and HEPES, buffered with KOH to pH: 7.4.

#### 2.8.2. Plasma membrane potential modulation

Cells were resuspended in HBSS at 5 × 10^5^/ml and placed into FACS tubes. Immediately before measurements, cells were incubated for 5 min using various concentrations of ZG613, solvent (DMSO) or KCl (50 mM) as indicated in the Results, following additional 5 min incubation with 100 nM Anionic Oxonol Dye DiBAC4(3) (D8189, Sigma Aldrich, MO, USA). Fluorescence was analysed on a BD FACSCalibur flow cytometer, and increase in red fluorescence was calculated in FlowJo (TreeStar, Inc.).

#### 2.8.3. Non-invasive microelectrode ion flux measurements (MIFE)

To analyse K^+^ and Na^+^ ion transport across the plasma membrane of MCF7 and HMLETwist cells, we used MIFE. MIFE system^64^ is based on ion flux estimation using ion selective microelectrodes that measure net ion fluxes (voltage gradient over the travel range) between two positions near the biological sample. Cells were seeded onto round glass coverslips placed in wells of 6-well plates (5 × 10^5^ cells in 350 μl medium), left for 15 min until cells are settled in the centre of each coverslip, and then the rest of the media was added in each well. HMLETwist cells were grown on coverslips coated with rat tail Collagen I (Gibco, MA, USA). The next day, after reaching confluent monolayer, coverslips were removed from the cell’s media to the Measuring solution into petri dish (Sarstedt TC Dish, Ø35 mm, Standard) immediately before the MIFE measurements. We used high Na^+^ Measuring solution (10 mM HEPES, 137 mM NaCl, 1 g/L glucose, 0.5 mM MgCl_2_, 1 mM CaCl_2_, 5 mM KCl) and low Na^+^ Measuring solution (10 mM HEPES, 137 mM Choline, 0.2 mM NaCl, 1 g/L glucose, 0.5 mM MgCl_2_, 1 mM CaCl_2_, 5 mM KCl). After placing the petri dish with the sample into MIFE^TM^ 3 system (University of Tasmania), measuring microelectrodes (K^+^ and Na^+^) were adjusted just close above the monolayer of the cells by MIFE custom-assembled Narishige manipulator system consisting of MHW-4 micromanipulator (5:1 movement) and SM-17 electrode holder (Narishige, Tokyo, Japan). Movement of electrodes was provided by MIFE-MDRIVE motor drive system (complete) for hydraulic micromanipulator^65^. Samples and electrodes were observed by inverted Zeiss PrimoVert microscope. Reference electrode (1 M NaCl, 2% agarose) was placed in Measuring solution on the edge of the petri dish. Vehicle (0.5% DMSO) or 2 μM and 10 μM ZG613 were added during each measurement after initial equilibration of cells (approx. 10 minutes) in Measuring solution. After addition of tested compounds, response signal of cells was recorded during 10 minutes of valid signal. Microelectrodes were directly connected to MIFE 3 preamplifier and the signal was then amplified and converted by MIFE 3 8-channel controller module, which was directly connected to PC by USB cable. System settings, data acquisition and basic flux calculation were performed by original MIFETM 3 software (MIFE, version 3.3.0.0.). Average ion fluxes were calculated using Excel.

#### 2.8.4. Intracellular Ca^2+^ measurements

Cells at 10^5^/ml were resuspended in DMEM without serum supplemented with 0.02% Pluronic F-127 (Sigma) and 3 μM Fluo-4 AM dye (BD Biocsiences). After 60 min incubation in the dark at 37 °C, cells were washed and incubated for further 20 min in the same medium without Pluronic. After additional washing, cells were resuspended in HBSS with or without calcium, and treated with DMSO or ZFG613 immediately before measurements. Fluorescence was analysed on a BD FACSCalibur flow cytometer, and increase in green fluorescence was calculated in FlowJo (TreeStar, Inc).

### 2.9. Tumour growth inhibition in vivo

#### 2.9.1. Animals

Female BALB/c 8 weeks old mice (20-25 g), bred at Ruđer Bošković Institute, were used. Animals were housed at a constant temperature (22 °C) and under a light cycle of 12 hr light/12 hr darkness (lights on at 7.00 a.m.). They were six animals per experimental group, 3 animals in each cage. Food and water were freely available. Prior to the experiment, the animals were not habituated to intraperitoneal (i.p.) injections. All animal care and experimental procedures were carried out in accordance with the Directive 2010/63/EU of the European Parliament and Council of the European Union on the protection of animals used for scientific purposes and the Croatian law on animal welfare. The ethical approval was obtained from Ruđer Bošković Bioethics Committee.

#### 2.9.2. Acute cytotoxicity

The acute cytotoxicity study of ZG613 was performed according to protocol: OECD AOT 425^66^ with modifications. Single animal tested received 45 mg/kg, s.c, dose that was lower than LD_50_ estimated *in silico* (600 mg/kg). The dose applied was the maximal dose available, prepared by dissolving ZG613 in preheated (60 °C) Corn oil (Sigma). Tested dose didn’t kill the animal, but it wasn’t increased due to the low solubility of ZG613. We daily administered the same dose during one week. After that, the experiment was terminated, since there were nor weight loss neither any other harm to the tested animal.

#### 2.9.3. Tumour growth inhibition

Tumours were induced in female animals by injections of murine breast cancer cells 4T1 in hind limbs, 10^6^ cells/100μl RPMI medium. After one week, we started with daily administration of ZG613 (45 mg/kg, i.p.) dissolved in Corn oil (ZG613 group), whereas animals from the control group were treated with Corn oil only. There were six animals per each experimental group at the beginning of the experiment, and 5 in the ZG613 group and 4 in the control group at the end of experiment. Each group was divided into two cages. Tumour size measured by a calliper and animal weights were monitored in 2-4 days intervals. Animals were sacrificed 18 days after the beginning of the therapy, and numbers of metastasis in lungs were counted under the magnifier.

### 2.10. Statistical Analysis

Results are represented as mean values from at least three separate experiments, unless stated differently. All graphics with error bars are presented as mean ± standard deviation (SD) and were generated in GraphPad Prism 5 software except for MTT curves where data was plotted using Excel. To determine statistical significance between samples, we used one-way ANOVA with post-hoc Tukey’s multiple comparison test except for *in vivo* analysis, membrane permeability and Ca^2+^ measurements where data were analysed using unpaired t test. Statistical calculations and generation of graphics were performed in GraphPad Prism (NS - non-significant; * - p < 0.05, **- p < 0.01, *** - p < 0.001).

## 3. Results

### 3.1. Molecular dynamics simulations of ZG613 behaviour in water/hydrophobic surfaces

First, to assess the ionic affinity, MD simulation was performed by solvation of compounds in water together with a variety of electrolytes. Potassium and sodium ions were used in order to investigate any competition between these two cations. During these MD simulations, multiple binding events between ZG613 and potassium ions can be observed, and molecules seem to be “locked” in the conformation adopted after K^+^ binding (video 1, Supporting information). However, it appeared that sodium ions were not able to form a complex with the crowns. Our findings corroborate the previously published experimental data. Based on the cavity size, 18-crown-6 compounds readily bind monovalent cations, with a higher binding constant (Ks) for K^+^ than for sodium ions^67^.

The simulations containing the water/chloroform interface were performed to assess the efficiency of ion transport. The compounds dissolved quickly within the chloroform phase, within the first couple of nanoseconds on average. While some binding events with potassium were observed in the water phase, crowns did not go inside the chloroform together with any ion. While a strong aggregation was noted in bulk water, once the compounds were dissolved within the chloroform, no cluster was observed (Fig. 2A; video 2, Supporting information).

**Figure 2.**
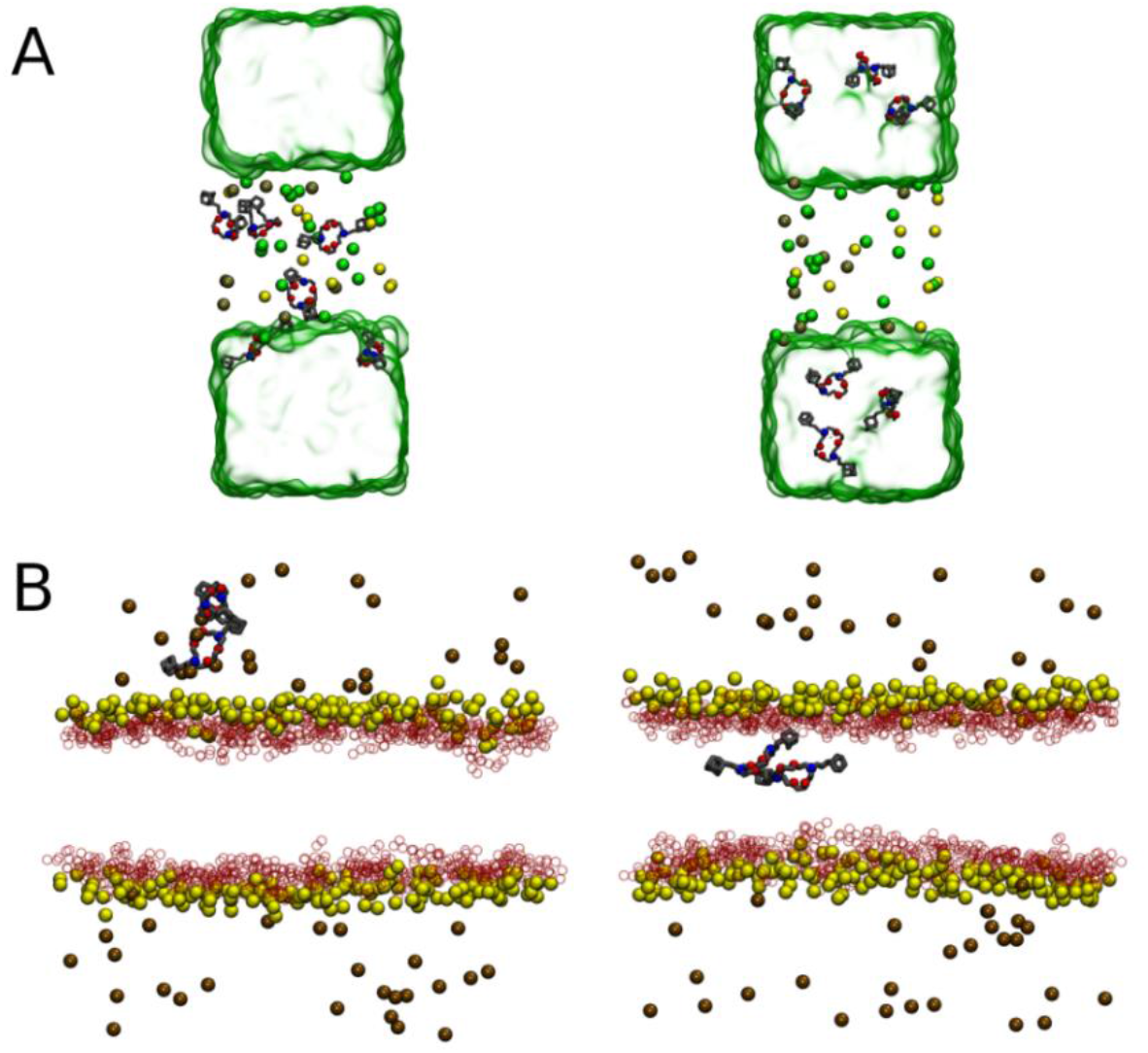
Molecular Dynamics (MD) simulation of ZG613 behaviour in water/chlorophorm interface and POPC membrane. Snapshots from biphasic system water/chloroform containing six crown ethers (A). Left column: starting configurations. Right column: representative snapshots of the equilibrated systems. The chloroform phase is depicted using a green glassy box. Ions are represented by coloured balls. Brown, yellow and green are used for potassium, sodium and chloride respectively. Crown ethers are in grey with the coordinating atoms, oxygen and nitrogen represented using red and blue balls respectively. Representative snapshots of the POPC membrane system with two ZG613 molecules (B). Left: Starting configuration. Right: Final configuration after a 100 ns molecular dynamics simulation. Phosphorus from the headgroups and oxygen atoms from the choline moieties are depicted with opaque yellow and transparent red balls respectively. Potassium ions are shown as brown balls. Crown ethers are in grey with the coordinating atoms, oxygen and nitrogen represented using red and blue balls respectively.

Additional simulations including a membrane constituted by phosphatidylcholine (POPC) molecules, a paragon for biological membrane, were performed to study ZG613 impact upon biological membranes. A direct diffusion of the compound from the water region towards the centre of the membrane was observed. Thanks to their functionalized moieties ZG613 molecules passes directly the headgroup region of the lipids and diffuses easily toward the centre of the membrane (Fig. 2B; video 3, Supporting information). The pristine diaza-18-crown-6 remained at the level of the ester region of the lipids (video 4, Supporting information). When the simulations contained 8 (video 5, Supporting information) and 4 molecules, ZG613 clustered together and remained within the bulk water during the whole simulation period. In the simulations with two ZG613 molecules, one of the molecules was seen to come toward the membrane interface bound with a potassium ion. However, upon entering the hydrophobic core of the bilayer, the ion dissociated and came back to the bulk water phase, while the compound continued its way toward the centre of the bilayer.

Molecular dynamics simulation indicates that ZG613 readily diffuses deeply into the centre of the lipid bilayer is able to bind K^+^ but is not able to transport cations through the membrane. In addition, ZG613 forms large aggregates that are orientated towards the water phase of the membrane.

### 3.2. ZG613 inhibits growth and induces death of breast cancer cells and EMT-model cells in vitro

We previously observed ZG613 cytotoxicity toward different human tumour cell lines^39^. In the prevailing study, we broaden the focus on cell lines that represent chemotherapy-refractory cancer and bear breast CSCs characteristics. We performed cytotoxicity screening *in vitro* using MTT assay in breast cancer cell lines MCF7 and SUM 159, breast EMT-model HMLETwist and HMLEpBp cells and human fibroblasts MJ90. MCF7 cells express estrogen receptor (ER) alpha, mimicking invasive human breast cancers that express ER, while SUM 159 represents human TNBC model enriched with CSCs^68^. HMLE-Twist represents breast EMT-model where immortalised breast primary epithelial cells HMLE are forced to pass EMT by overexpression of transcription factor Twist^24^. Human fibroblasts MJ90 were included in the experiments as a non-cancerous cell line. Cells were treated with serial dilutions of ZG613, and percentage of growth and IC_50_ values were analysed after 72 hr according to drug screening platform previously established in our lab. In addition, we performed shorter alternative of MTT assay to determine the beginning of cell death. ZG613 showed cytotoxic activity toward all of the cell lines, with IC_50_ values between 0.6 μM - 3 μM (Fig. 3A). TNBC cell line, SUM159, was slightly but not significantly more sensitive comparing to MCF7 cells. SUM159 were significantly more sensitive to ZG613 comparing to fibroblasts MJ90 (Fig. 3A).

**Figure 3.**
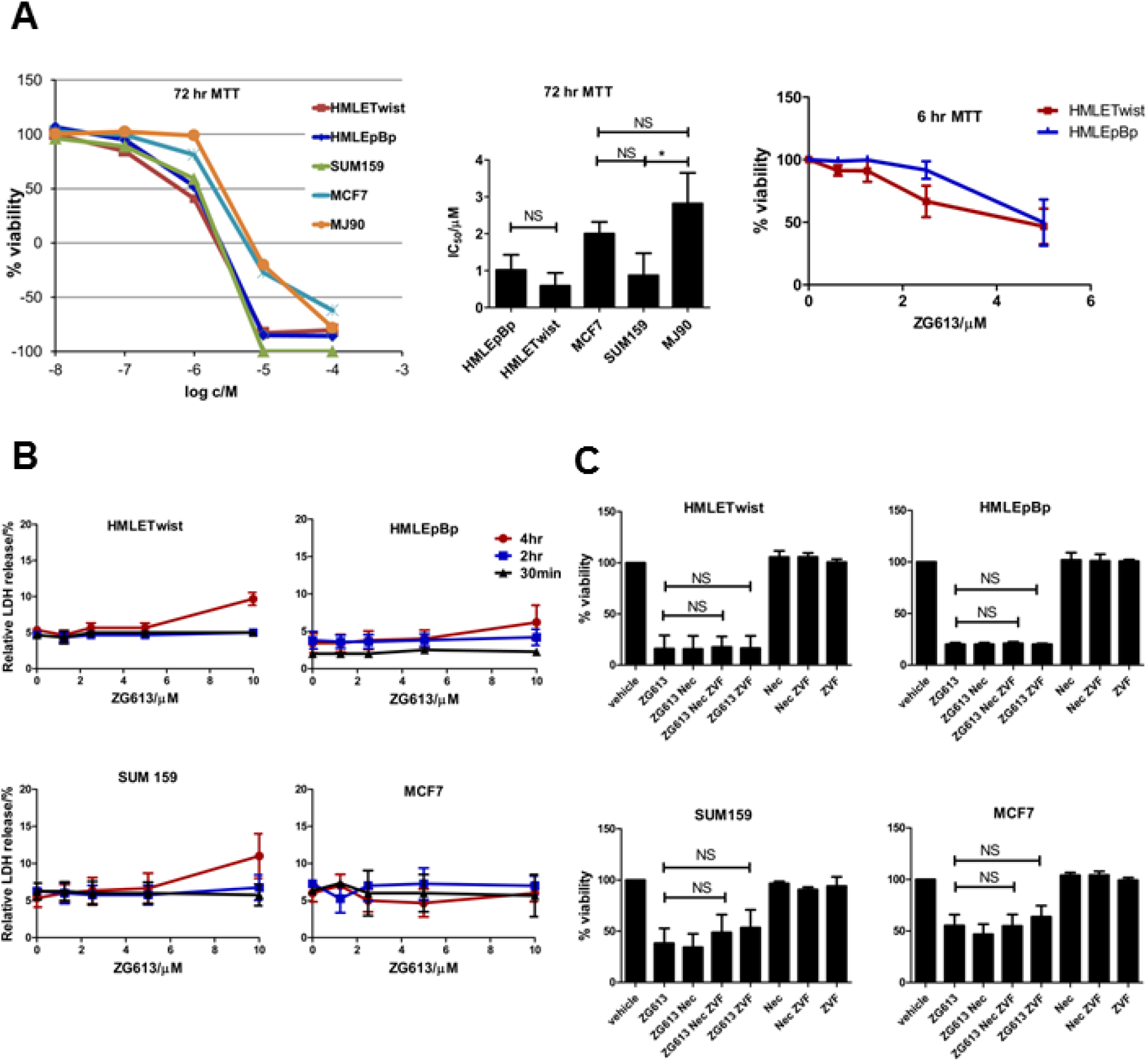
ZG613 reduces viability and induces necrotic cell death. Dose response curves and IC50 values of tested cell lines (A). Cytotoxicity of ZG613 was assessed using MTT assay 72 hr or 6 hr after the addition of ZG613. Curves and histograms present data from at least three independent experiments. Data were analysed using 1 way ANOVA with Tukey’s Multiple comparison test (*p < 0.05, NS-non-significant). To perform LDH release assay, cells were treated for 30 min, 2 and 4 hr with vehicle (DMSO) or various concentrations of ZG613 (B). Plots present data from at least three independent experiments. LDH release values were normalised to LDH positive control provided by the manufacturer. Histograms (C) represent growth inhibition of SUM159, MCF7 and HMLETwist/pBp cell lines by 5 μM ZG613 alone or in the combination with necroptosis inhibitor Necrostatin (Nec-1) and apoptosis inhibitor Z-VAD-FMK (ZVF). MTT assay was performed 24 hr after the treatment with vehicle (DMSO), inhibitors and 5 μM ZG613. Histograms represent data from at least three independent experiments. Data were analysed using 1 way ANOVA with Tukey’s Multiple comparison test (NS-non-significant).

In HMLETwist/HMLEpBp model, we observed low but not significant increase of ZG613 toxicity toward EMT cells HMLETwist compared to HMLEpBp, based on 72 hr treatment. To address possible early differential response, we performed 6 hr MTT assay, where EMT cells in 2.5 μM concentration were more sensitive than epithelial control (Fig. 3A). In general, ZG613 treatment resulted in a very steep dose-response curve in all cells, especially in the concentration range between 1-10 μM.

In order to separate rapid effects on cell membranes from the total membrane lysis that results from the death cascade, we aimed to pinpoint the beginning of cell death induced by ZG613 using LDH release assay. Release of LDH and other cytoplasmic components into the cell’s surrounding indicates severe membrane lysis and subsequent cell death. Cells did not start to die until 4 hr after the treatment in all tested cell lines (Fig. 3B). When lower concentrations were used, longer time was necessary for induction of cell death. Successful cytotoxic effect is proportional to the time and concentration, pointing to a conclusion that cells seem to resist the compound.

We aimed to distinguish modality of cell death between apoptosis/necroptosis/necrosis using inhibitor of necroptosis Necrostatin 1 (Nec-1) and inhibitor of apoptosis Z-VAD-FMK (ZVF) according to previously described procedure^69,70^. Nec-1 and/or combination of Nec-1 and ZVF was used to inhibit necroptosis, ZVF was used to inhibit apoptosis, and modified MTT assay with inhibitors was performed during 24 hr. Small but not significant rescue of MCF7 and SUM159 cells was observed using inhibitor of apoptosis (Fig. 3C). In all of the tested cell lines, death induction by ZG613 was not significantly reversed using inhibitors, indicating that cells don’t die via these two modalities of regulated cell death.

### 3.3. ZG613 induces differential expression of genes in EMT-model cells

To identify molecular targets in cells, we analysed transcriptome of HMLETwist and HMLEpBp cells 24 hr after the treatment with 1 μM ZG613. Duration and concentration was chosen in order to accumulate the majority of signalling responses while at the same time to avoid triggering of the cell death cascade. We performed RNA-Seq using Illumina platform. Significantly upregulated and downregulated genes relative to control (vehicle-treated cells) for each cell line are presented in Figure 4.

**Figure 4.**
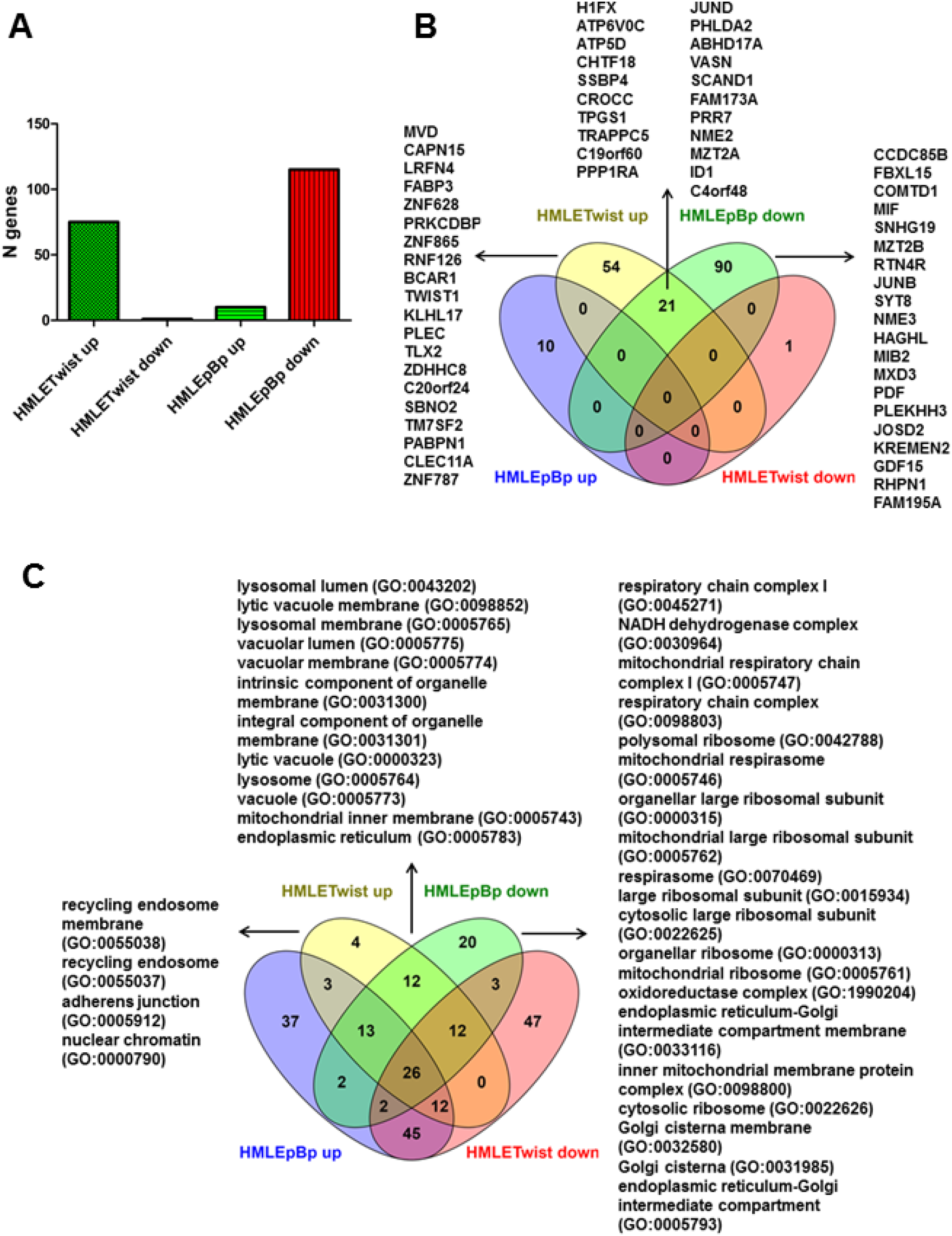
ZG613 induces differential expression of genes in HMLETwist/HMLEpBp model. Histogram A represents significantly up-regulated and down-regulated genes among differentially expressed gens in HMLETwist and HMLEpBp cells induced by ZG613. (B) Presentation of significant differentially expressed genes by Venn diagrams. Diagram contains 4 datasets that present significant differentially expressed genes in HMLETwist and HMLEpBp cell lines after the treatment with ZG613. First 20 genes exclusively up-regulated in HMLETwist and downregulated in HMLEpBp cell line are listed. (C) Venn diagram of 4 datasets that present Cellular compartment GOs related to differentially expressed genes in HMLETwist and HMLEpBp cell lines after the treatment with ZG613. All GOs exclusively up-regulated in HMLETwist and downregulated in HMLEpBp cell line, as well as overlapping GOs from the same two datasets are listed.

Significant differentially expressed genes are presented using Venn diagrams in Figure 4B, where clear segregation between EMT (HMLETwist) and epithelial control (HMLEpBp) cell lines is evident. First 20 genes, based on log2(fold change) ranking, representing exclusively downregulated dataset in HMLEpBp and exclusively upregulated dataset in HMLETwist, are listed (Fig. 4B). Genes involved in membrane lipid composition (MVD, FABP3, PRKCDBP, ZDHHC8, TM7SF2) are predominantly represented among first 20 in HMLETwist-up dataset, and additionaly represented in HMLEpBp–up (ACAT2) and overlapped HMLETwist-up/HMLEpBp-down (SCAND1) datasets (Fig. 4B). Furthermore, among first 20 genes in each dataset, there are genes involved in vesicular transport (RNF126, TRAPP, ATP6V0C, KREMEN, RAB3B, SYT8), cytoskeletal remodelling (ABL2, LRFN4, KLHL17, PLEC, TPGS1, CROCC, RTN4R, RHPN1) and mitochondrial functions (PABPN1, FAM153A, ATP5D, PDF), equally distributed among datasets. Other genes listed among first 20 in each dataset, if functions are determined so far, are involved in diverse aspects of proliferation, differentiation and transformation-related signalling (CAPN15, BCAR1, TWIST1, CLEC11A, IGFBP3, AKAP12, ALDH1A3, MIB2, MXDR, PHLDA2, JUND, VASN, NME2, GDF15, ID1). ZG613 downregulates genes involved in inflammation (MIF and CXCL8) exclusively in epithelial cells. Single significantly downregulated gene in EMT cells is KRT6, gene characterised as the promoter of EMT. Lists of all significant differentially expressed genes are shown in Table S1 (Table S1, Supporting information). Furthermore, all up and down-regulated genes with fold change above 1.1 were included into Gene Ontology (GO) analysis using Statistical overrepresentation test available on Panther web platform. Venn diagrams representing Cellular compartment GOs are presented in the Figure 4C, with lists of exclusively upregulated GOs in HMLETwist, downregulated GOs in HMLEpBp, and overlapping GOs from these two datasets. We didn’t consider HMLETwist-downregulated GOs since there was only one significantly downregulated gene in that dataset. Cellular compartment GO analysis pointed to upregulation of genes involved in vesicular trafficking in EMT cells, while the same processes are downregulated in epithelial controls. Additionally, mitochondria and Golgi associated processes are exclusively related to downregulated genes in epithelial cells. In summary, ZG613 affects expression of genes involved in membrane lipid composition, vesicular transport and cytoskeletal remodelling, all of which is crucial for maintenance of membrane integrity. Genes involved in membrane lipid composition are more affected in EMT cell line. Moreover, GO analysis pointed toward plasma membrane recycling in EMT cells, while in epithelial cells treatment profoundly affected mitochondrial functions.

### 3.4. Membrane permeability changes

Since MD simulation suggests that crown ether ZG613 diffuses deeply into the lipid bilayer, while RNA-Seq indicates induction of processes related to maintenance of membrane integrity, we tested ZG613 ability to physically disrupt plasma membrane by induction of small ruptures that will cause increase in membrane permeability^71^. We quantified the influx of SYTOX Green dye into the cells during 2 hr incubation with 10 μM ZG613 (Fig. 5A). ZG613 increases general membrane permeability in all cells during 2 hr, while LDH leakage was not detected in any of tested cell. Membrane permeability remained intact when smaller concentration (1 μM) was used (Figure S1, Supporting information).

**Figure 5.**
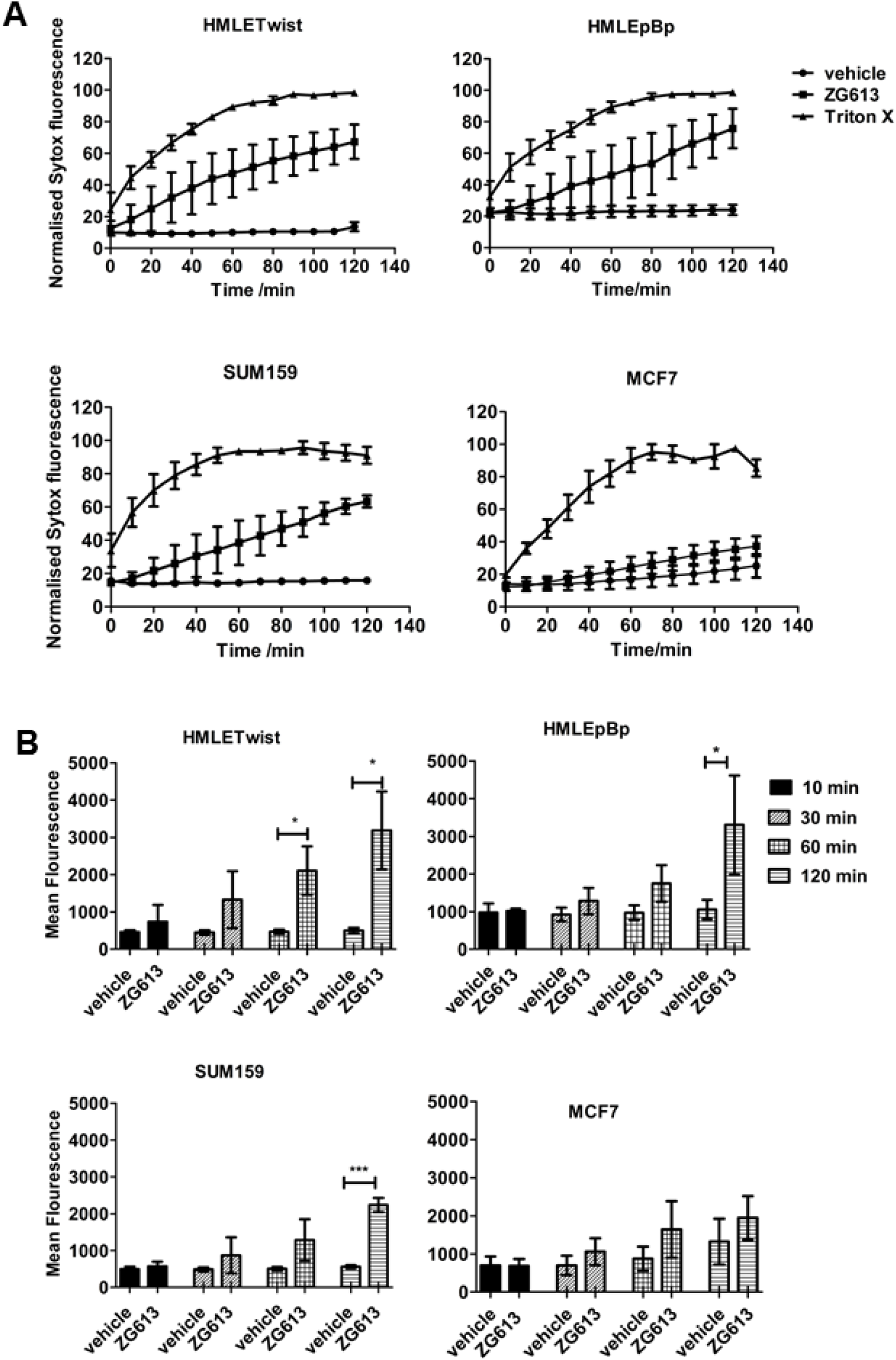
Study of membrane permeability induction by ZG613. Time course of Sytox influx induced by ZG613 is presented on panel A. Permeation of Sytox dye was monitored in breast cancer and breast EMT-model cell lines up to 120 min after the addition of 10 μM ZG613, DMSO or Triton X. Fluorescence was measured every 10 min using Tecan M200 microplate reader. Plots present data from at least three independent experiments. Maximal value obtained with non-ionic detergent Triton X (0.3 %) was set as 100%. (B) Membrane permeability changes at selected time points after ZG613 addition. Histograms present Sytox influx in 10 min, 30 min, 60 min and 120 min time points. Mean fluorescence values of vehicle-treated control (0.5% DMSO) and cells treated with 10 μM ZG613 were plotted. Sytox influx compared to vehicle-treated control in each time point is analysed using unpaired t-test, and significant Sytox influx is presented using asterisks (* p<0.05, *** p<0.005).

HMLETwist cell line was the most prone to permeability changes by the compound, where Sytox started to permeate in 10 min (Fig. 5B). In other cell lines, influx was noticeable after 30 min. Significant permeability increase was measured in 60 min in HMLETwist while in 120 min in HMLEpBp and SUM159 cell line. In cell line MCF7 we obtained non-conclusive data regarding ZG613 effects. Cells seem to be resistant to the permeability changes, but on the other hand, permeability was also increased by DMSO.

### 3.5. Mitochondrial function impairments by ZG613

Mitochondria are membrane bounded organelles that possess changed structure and function in tumour and EMT, as mentioned in Introduction. Since mitochondria possess more negative membrane potential than their surroundings in eukaryotic cell^72^, potential cation-binding ionophoric drugs, like ZG613, are expected to have an impact. Moreover, our RNASeq data pointed toward mitochondria as potential ZG613 targets. Hence, we assessed their vulnerability to membrane perturbations by ZG613. To measure changes in mitochondrial transmembrane potential (ΔΨm), we used fluorescent lipophilic cationic dye JC-1. If membrane is compromised and membrane potential becomes more positive, JC-1 emission spectra shift from red to green. We observed mitochondrial membrane depolarisation using flow cytometry 60 min after the addition of ZG613 in all tested cell lines (Fig. 6A). Uncoupler of mitochondrial oxidative phosphorylation FCCP (10μM) was used as a positive control.

**Figure 6.**
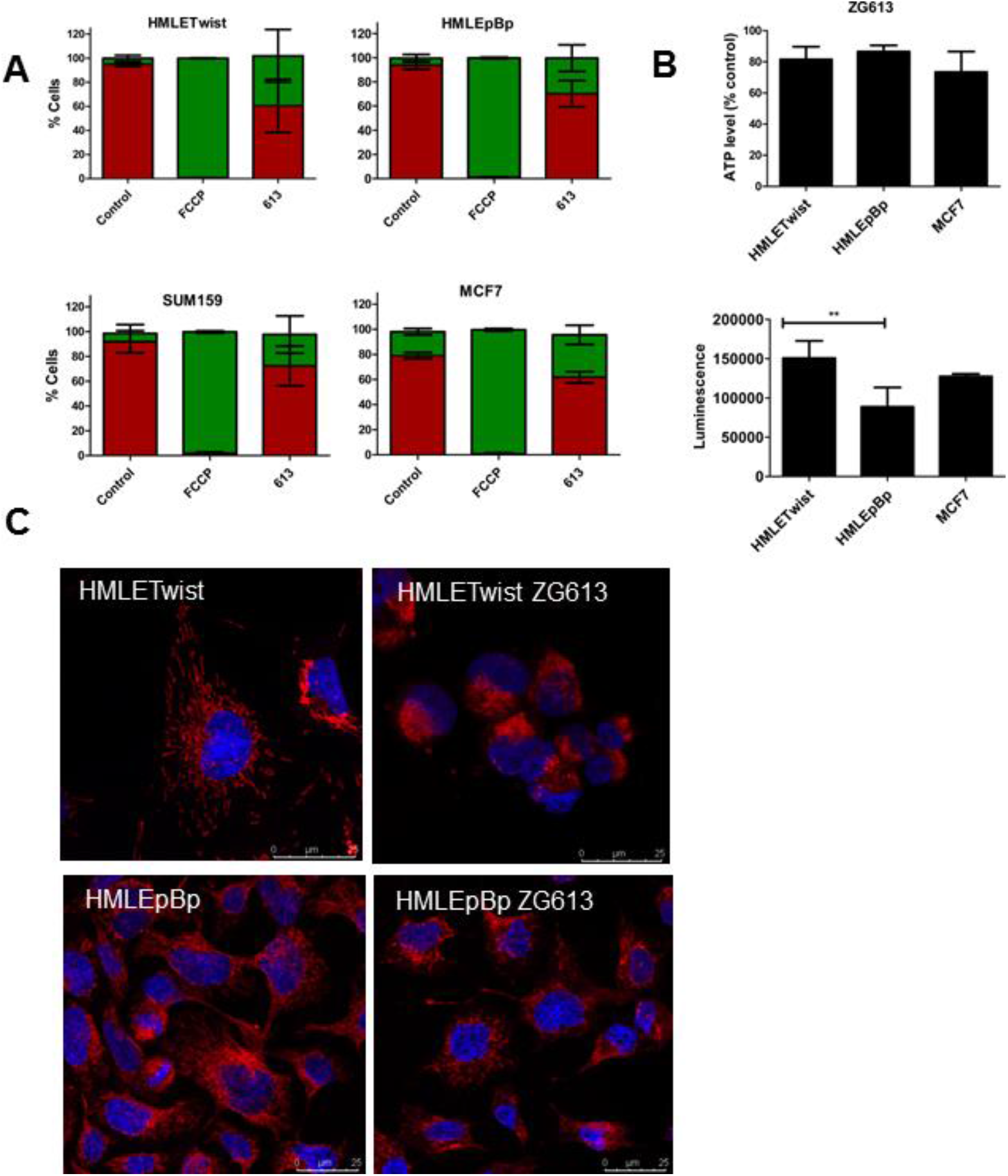
Mitochondrial function impairments by ZG613. (A) ZG613 depolarises mitochondrial membrane potential. Histograms represent flow cytometry analysis of mitochondrial membrane depolarisation in breast cancer and breast EMT cells using JC-1 cationic dye. Cells were treated with 10 μM ZG613 or 10μM FCCP for 60 min. Histogram (B; up) represents relative ATP levels upon ZG613 treatment in HMLEpBp, HMLETwist and MCF7 cell lines normalised to vehicle-treated control (0.5 % DMSO). Cells were treated with 10 μM ZG613 for 30 min, and relative ATP levels were determined using CellTiter-Glow assay. Histogram (B; down) represents ATP levels in vehicle-treated controls compared among different cell lines. Histograms represent data from at least three independent experiments. Data were analysed using 1 way ANOVA with Tukey’s Multiple comparison test (** p<0.01). (C) Changes in mitochondrial morphology by ZG613. Picture represents HMLETwist and HMLEpBp cells stained with Mitotracker DEEP RED FM, dye that visualises mitochondria (in red). Nuclei were stained with DAPI (in blue). Cells were treated with 5 μM ZG613 during 24 hr. Photographs were taken using Leica SP8 confocal microscope. Typical photographs of two independent experiments were shown.

Since it is shown by others that EMT cells acquire changes in mitochondrial morphology^34^, we investigated morphology of mitochondria in our EMT model. We used Mitotracker DEEP RED FM dye to visualise mitochondria and analysed cells using confocal microscopy. HMLETwist cells exhibit changed mitochondrial morphology when compared to isogenic control HMLEpBp. Although we couldn’t specify the nature of the changes observed, mitochondrial appearance resembles mitochondrial hyperfusion. Hyperfusion is an EMT related feature associated with activation of mitochondrial fusion protein 1 (MFN1) and elevation of ATP levels^34^. Interestingly, treatment with 5 μM ZG613 changed mitochondrial morphology in HMLETwist, but not in HMLEpBp (Fig. 6C).

The primary function of mitochondria is generation of ATP, therefore we investigated cellular ATP content shortly after addition of 10 μM ZG613. Moderately decreased cellular ATP levels (between 14-27 %) 30 min after the addition in all of the cell lines tested were observed (Fig. 6B). We used CellTiter-Glo luminescent assay to measure relative ATP levels. Moreover, in consistence with mitochondrial morphology experiments, HMLETwist cells possess increased levels of ATP compared to HMLEpBp cells.

### 3.6. Ion transport study

MD simulation, as well as our previous study of ion transport, gave us adverse answers regarding ZG613 potential ionophoric properties. Therefore we performed an analysis of ZG613 impact on the ion homeostasis. At the beginning of the study, we performed quantitative measurements of resting membrane potential (Vm) in HMLETwist, HMLEpBp, SUM159 and MCF7 cell lines using patch clamp/current clamp technique (Fig. 7C). Resting membrane potential is an intrinsic property of any cell that is related to the differentiation/stemness capacity, in a way that progenitor cells have more positive values than differentiated progeny, all of which could be manipulated using ionophoric compounds^73^. We observed that both HMLETwist and HMLEpBp have a rather positive Vm value (around - 20 mV) comparing to HMEC cells from which they originated, and which have around - 60 mV^74^. Unfortunately, observed disturbance in bioelectricity occurs nonexclusively in all HMLE cells, unabling investigations of potential EMT-specific bioelectricity. MCF7 cell line had the most negative Vm in our experiments.

**Figure 7.**
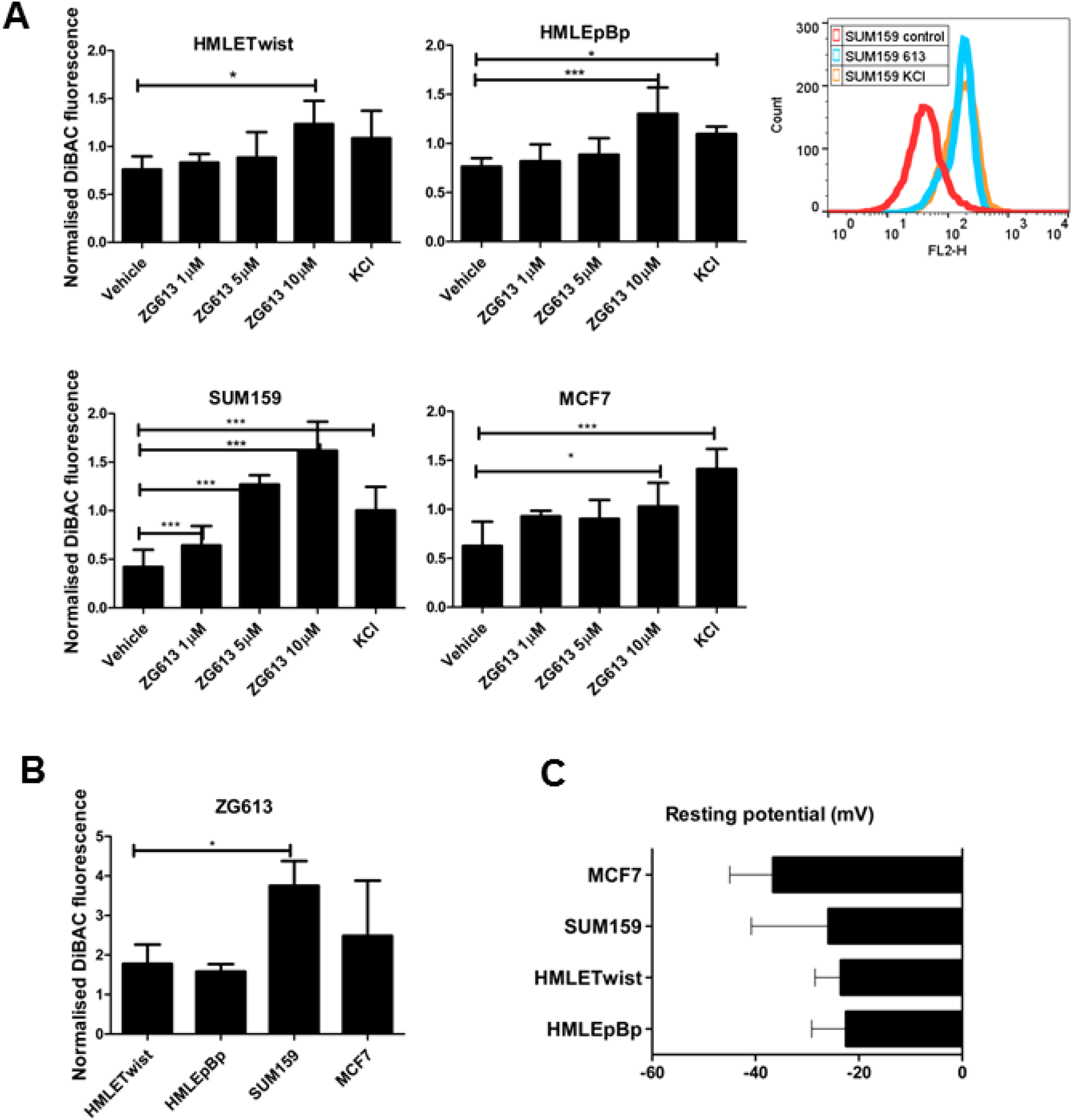
Measurements of plasma membrane potential modulations by ZG613. (A) Measurements of membrane potential upon ZG613 addition. Histograms represent flow cytometry analysis of cells stained with DiBAC4(3) and treated with various concentrations of ZG613 for 10 min. Control cells were treated with vehicle (DMSO), and KCl was used as a positive control that depolarises cells. Histograms represent normalised median fluorescence values from at least three independent experiments normalised to control in each separate experiment. Data were analysed using 1 way ANOVA with Tukey’s Multiple comparison test (* - p < 0.05, ** - p < 0.01, *** - p< 0.001). Representative flow cytometry histogram shows shifts in fluorescence upon ZG613 and KCl treatment in SUM159 cells. (B) Comparison of depolarisation levels among cell lines. In each cell line, values are normalised to vehicle-treated control. Data were analysed using 1 way ANOVA with Tukey’s Multiple comparison test (* - p < 0.05). (C) Nystatin-perforated patch clamp was performed to measure resting membrane potential in breast cancer and EMT-model cells. Histograms represent at least 6 independent recordings per each cell line.

Furthermore, we measured changes in the membrane potential of cells rapidly after the treatment with the compound. We applied DiBAC4(3) anionic dye that accumulates in cells if membrane potential become more positive*, e.g*. if cells depolarise, and performed flow cytometry. Control cells were treated with vehicle (DMSO), while KCl was used as a positive control. ZG613 significantly depolarises all of the cell lines in 10 min in a dose-dependent manner (Fig. 7A). Since general membrane permeability was increased only in HMLETwist at this time point (Fig. 5B), we presumed that depolarisation observed is a consequence of changes in ion fluxes in HMLEpBp, MCF7 and SUM159 cells. Cell line SUM159 was the most sensitive to depolarisation by the compound (Fig. 7B), while there was no difference between HMLETwist and HMLEpBp cells.

We performed MIFE analysis in HMLETwist and MCF7 cells to evaluate direct impacts of ZG613 on sodium and potassium ion fluxes. We added non-ionic detergent Digitonin (12.5 μg/ml) at the end of each measurement, to assure quality of each measurement by inspection of both levels and direction of fluxes. Influx is presented using positive, while efflux using negative values. Digitonin induced robust Na^+^ influx from medium into the cells, and K^+^ efflux from cells into the medium in physiological conditions (Fig. 8A), *e.g.* conditions when measuring solution consisted of high sodium and low potassium ion concentrations.

**Figure 8.**
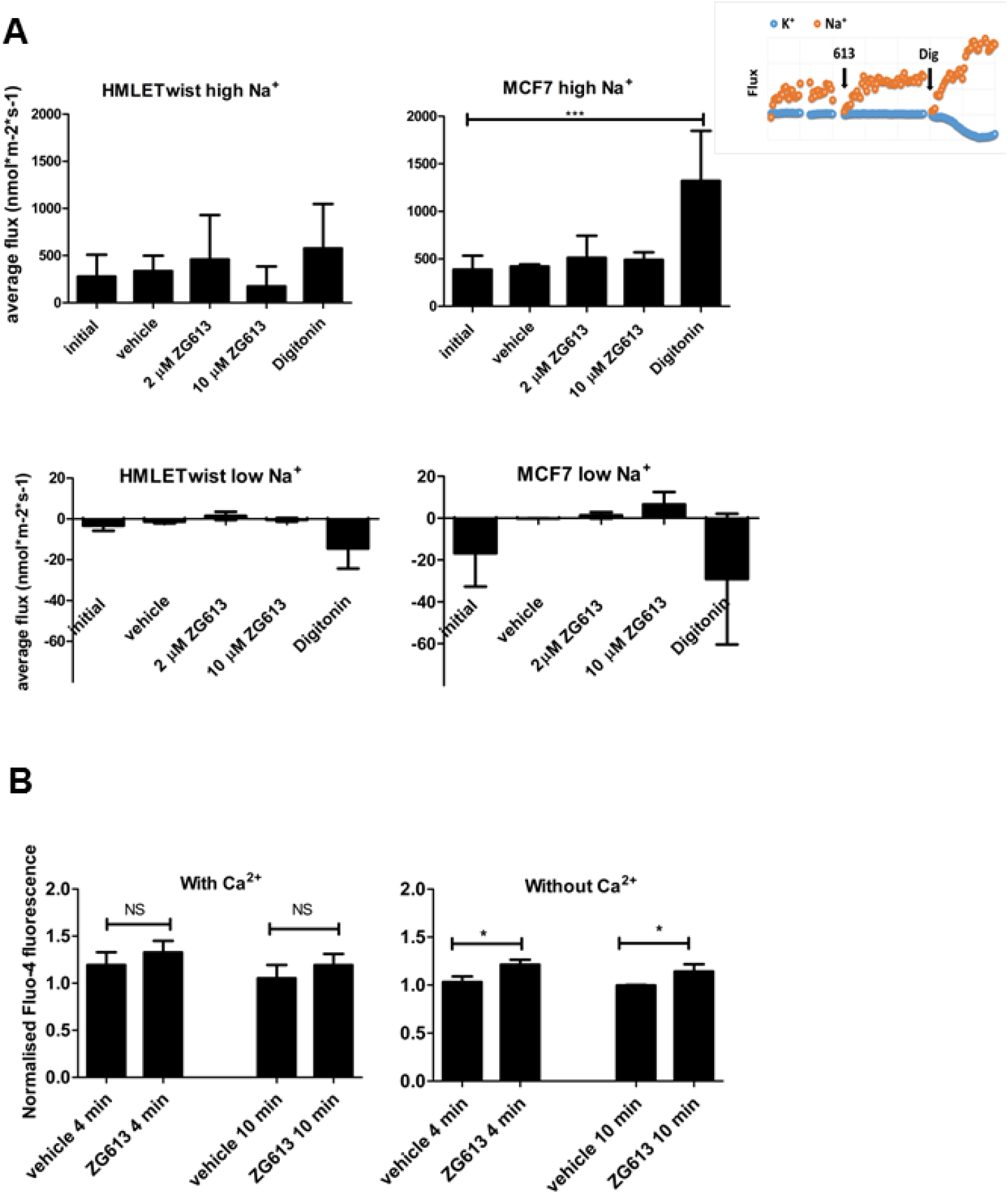
Measurements of ion fluxes. (A) Histograms represent average fluxes of Na^+^ induced by ZG613, vehicle (DMSO, 0.5%) and non-ionic detergent Digitonin (12.5 μg/ml) in HMLETwist and MCF7 cell lines. High Na^+^: measuring solution with physiological Na^+^ concentration (137mM). Low Na^+^: measuring solution with low Na^+^ concentration (0.2 mM). Data were analysed using 1 way ANOVA with Tukey’s Multiple comparison test; *** p<0.005. Typical recording of K^+^ and Na^+^ fluxes in MCF7 cells treated with ZG613 (613) and Digitonin (Dig) is plotted on the right. (B) Measurements of intracellular Ca^2+^ using Fluo-4 dye. MCF7 cells were incubated with calcium selective fluorescent dye Fluo-4 for 30 min and treated with DMSO or 10 μM ZG613 for 4 and 10 min. Changes in fluorescence were measured using the flow cytometer. Histograms represent data from at least three independent experiments. Median fluorescence intensity values are normalised to non-treated controls. With Ca^2+^ : HBSS with calcium; Without Ca^2+^ : HBSS without calcium. Data were analysed using unpaired t test (NS – non-significant; * - p < 0.05).

In MCF7 cells, both low (2 μM) and high concentration (10 μM) of ZG613 induces minuscule Na^+^ influx that was measured during 10 min after the addition of the compound (Fig. 8A). To substantiate results, we performed the same analysis using low Na^+^ measuring solution (non-physiological conditions). Interestingly, in low Na^+^ conditions, we expected efflux of sodium ions from cells to the medium upon ZG613 treatment, as induced by Digitonin, but we observed inversion-small influx of sodium ions into the cells. In HMLETwist cells, we obtained non-conclusive data, since in these cells fluctuations of fluxes were large already from the beginning in each measurement. Nonetheless, in low Na^+^ measuring solution, the effect of the compound on Na^+^ flux was negligible. We didn’t observe induction of potassium ion fluxes in any of the cell line tested after the compound addition (Figure S2, Supporting information). In general, ZG613 induced miniscule, statistically insignificant sodium ion influx in MCF7 cells using both physiological and low Na^+^ conditions during 10 min, prior to the increase in general membrane permeability. Although we proposed the mechanistic capability of the compound to increase membrane permeability for Na^+^, changes in sodium ion fluxes induced were too small and therefore could be neglected.

Since ZG613 depolarised cells, but barely induced Na^+^ and K^+^ fluxes, we investigated whether there is an increase in intracellular Ca^2+^ shortly after the ZG613 addition. Changes in intracellular Ca^2+^ levels in MCF7 cell line were monitored using Ca^2+^ indicator, fluorescent dye Fluo-4, in medium with or without Ca^2+^ using the flow cytometer. Measurements in medium without Ca^2+^ were included in order to investigate whether the compound is able to induce Ca^2+^ depletion from endoplasmatic reticulum (ER) stores. We measured mild intracellular Ca^2+^ level increase 4 min after the addition of ZG613 in both media (Fig. 8B).

### 3.7. Effects on the tumour growth in vivo

The effects of ZG613 on the tumour growth *in vivo* were assessed using the single, non-toxic dose (45 mg/kg i.p.,) as shown in the Figure 9. Tumours were induced in female mice by injections of 4T1 cells in hind limbs. ZG613 was administered daily (45 mg/kg i.p.), while tumour size and animal weight were measured in 2-4 days intervals. We observed a mild retardation of tumour growth in ZG613 treated group, followed by a mild, but non-significant decrease of metastasis formation in lungs. The animal weight did not oscillate between groups, indicating that the compound wasn’t toxic. Since the compound was dissolved in corn oil, control group was administered with solvent.

**Figure 9.**
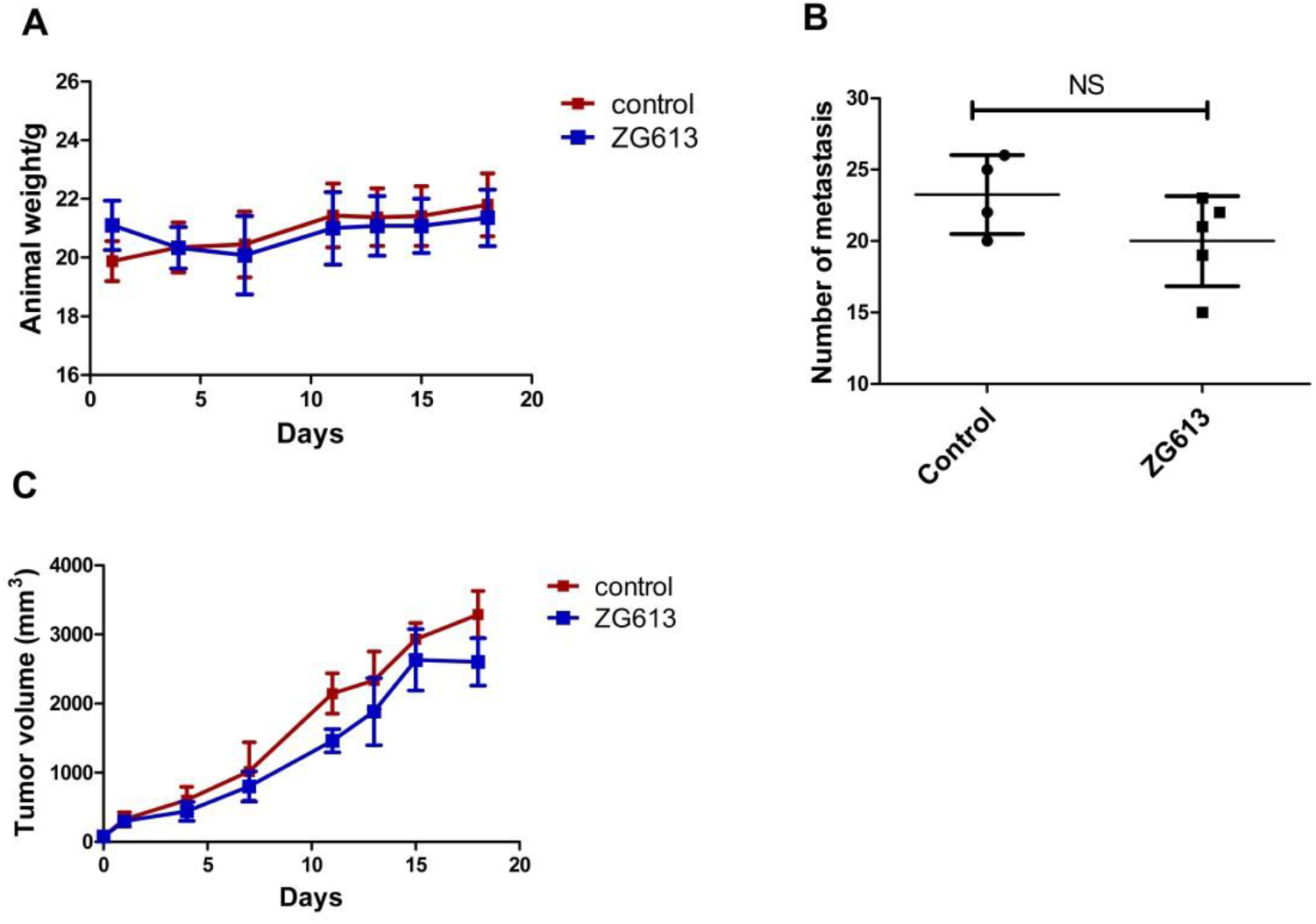
Effects of ZG613 on the animal weight, tumour growth and lung metastasis formation in mice. Tumours were induced in female animals by injections of 4T1 cells in hind limbs. ZG613 was administered daily (45 mg/kg i.p.), while tumour size (A) and animal weight (B) were measured in 2-4 days intervals. Animals were sacrificed 18 days after the beginning of the therapy, and numbers of metastasis in lungs were counted under the magnifier (C). There were 6 animals in each group at the beginning of the experiment, and 5 in the ZG613 group and 4 in the control group at the end of experiment. Data were analysed using unpaired t test; NS-non significant.

## Discussion

The present study clearly shows that diaza-crown ether ZG613 exhibits heterogeneous anticancer activity that hits multiple targets within the cell.

Based on the lack of interactions of 18-crown-6 derivatives with DNA, induction of DNA damage based cytotoxicity had been excluded in a previous study^38^. In addition, we did not measure significant induction of ROS nor changes in the cell cycle after the treatment that could point toward induction of DNA damage cascade (unpublished results). Thus, we tested the working hypothesis of physical disruption of plasma membrane, or/and disturbance in ion transport homeostasis as the basis of ZG613 antitumor activity, as described in the Introduction.

MD simulations revealed that ZG613 readily diffuses deeply into the POPC membrane (Fig. 2B). When more molecules were applied simulating higher concentration, ZG613 formed aggregates orientated toward the water side of the membrane. According to simulations, *e.g*. in water, water/chloroform and POPC membrane, the compound is able to bind K^+^ but not to transport it through the membrane, supporting data from Šumanovac et al^40^ that showed that ZG613 extraction ability is superior, but transport rates for both K^+^ and Na^+^ were low.

Compound ZG613 was highly cytotoxic in the concentration range between 1-10 μM (Fig. 3A). The shapes of dose response curves indicate that little change in concentration is needed to cause an increase in cytotoxicity. Careful selection of doses for each cell line in order to induce a robust effect without killing the majority of the cells was required. Unfortunately, this pointed toward an obstacle in potential use of ZG613 as an antitumor drug. Moreover, the length of treatment is also critical, indicating that there is point of no return for every cell line. When shorter time intervals are used cells suffer, but eventually recover. For example, when ~ 1 μM doses are applied for 6 hr, cells will retain ~ 90% viability, while the same concentrations have more severe impact after 72 hr (Fig. 3A).

Physical disruption of the plasma membrane is usually followed by a subsequent cell death via necrosis, necroptosis or apoptosis^75^. Bacterial pore-forming toxins induce necroptosis^76^, while dysregulation of ion homeostasis leads to induction of apoptosis, necrosis and necroptosis^6^. Available inhibitors of necroptosis and apoptosis did not have any major effect on the cell death induction by ZG613, indicating that cells don’t die via these two modality of regulated cell death (RCD), hence cells die most likely through necrosis (Fig. 3C). Nevertheless, observed effects of ZG613 on the plasma membrane integrity resemble patterns described by others, for example, to the effects of bacterial pore-forming toxins. Pore-forming toxins can also induce pyroptosis, another form of regulated necrosis, in immune cells^77^. Recently, it was shown that small molecules can trigger pyroptosis in breast tumour cells^78^. In addition, cationic amphiphilic drugs could be repurposed to induce lysosomal cell death in cancer^20^. Pyroptosis and lysosomal cell death overlap or culminate with non-regulated necrotic cell death and therefore couldn’t be absolutely excluded. The beginning of cell death starts after 4 hr according to LDH release (Fig. 3B), which allowed us to separate rapid effects on cells as depolarisation and increased permeability that occur prior to the cell death, from the membrane loss of function due to the membrane lysis.

Both EMT and control HMLE cells respond to treatment by changes in expression of genes involved in maintenance of membrane integrity as genes that contribute to lipid composition, cytoskeletal rearrangements and vesicular transport (Fig. 4B). Our data resemble studies using bacterial pore-forming toxins^79^, that revealed that plasma membrane repair after pore formation depends on enhancement of membrane fluidity and facilitated release of extracellular vesicles containing damaged membranes via calcium-activated lipid scramblase TMEM16F. Expression of genes after ZG613 treatment evidently differs between EMT and epithelial control cells.

Increase in plasma membrane permeability by 10 μM ZG613, before release of LDH, indicates formation of small ruptures that are sufficient to allow fluorescent reporter transport, but not sufficient to induce total loss of membrane integrity (Fig. 5A). Plasma membrane permeability can be typically restored in seconds after the minor breach^1^. However, when larger breaches are involved, spontaneous resealing is replaced with repair mechanisms involving vesicle-vesicle or vesicle-membrane sealing, related to calcium signalling^2^. We speculate that same mechanisms apply to membrane disturbance by ZG613. When added in a low concentration (1μM), ZG613 did not show any impact on permeability (Fig. S1), possibly by sufficient spontaneous resealing of the induced ruptures. Membrane permeability increases in time (Fig. 5A), and eventually leads to a total membrane loss of function, measured by LDH release (Fig. 3B). Since there is limited data available on pore formation mechanisms using small molecules, the exact mechanism of ZG613 action remains obscured. Even regarding the known amphiphilic drugs that target membranes, e.g. statins, the exact membrane-disruption mechanism is, at present, not fully elucidated^11^. Up to date, pore formation is mostly studied using acellular models as liposomes^80^, and in research using bacterial pore-forming toxines^76,79^. ZG613 has not changed cell volume, which indicated that osmolarity was maintained, probably by simultaneous leakage of ions, small molecules and proteins. EMT cells show enhanced potential for the permeabilization compared to isogenic control cells (Fig. 5B), possibly due to a reduction in plasma membrane stiffness and remodelling of cytoskeletal architecture, making them particularly prone to physical stress and instability caused by membrane disturbance.

We showed that ZG613 affects mitochondria, a membrane bounded organelle. Ionophore salinomycin was shown to impact mitochondria selectively in cancer cells^81^. We determined that ZG613 doesn’t affect Na^+^ and K^+^ transport through the plasma membrane (Fig. 8A, Fig S2). Hence, measured depolarisation of mitochondrial membrane potential (Fig. 6A) could be a consequence of ion dysregulation, but also a consequence of impairments of other mitochondrial functions that would lead to a loss of mitochondrial membrane integrity. Demonstrated ATP leakage from the tested cells 30 min after the addition of ZG613 (Fig. 6B) also indicates mitochondrial loss of function, but an increase in plasma membrane permeability may also contribute. Comparison of mitochondrial morphology between EMT and control epithelial cells (Fig. 6C), revealed alterations in EMT cells that resemble mitochondrial hyperfusion. This EMT related feature acquires activation of mitochondrial fusion protein 1 (MFN1) and elevation of ATP levels^34^. In fact we have shown in RNASeq experiments^82^ that HMLETwist overexpresses MFN1 compared to HMLEpBp. This is in line with the increased level of cellular ATP in EMT cells compared to isogenic control (Fig. 6B). ZG613 altered mitochondrial morphology exclusively in EMT cells, which could be used for the EMT specific targeting. Interestingly, RNAseq GO analysis (Fig. 4C) pointed to expression changes of mitochondria-related genes in epithelial cells. We could speculate that EMT cells may have a more robust mitochondrial system both structurally and functionally, that ZG613 somehow targets, but not on the expression level. Mitochondria represent a novel target in the contexts of drug-resistant cancers, which just recently draw more attention and should be further investigated.

We observed depolarisation of the plasma membrane 10 min after ZG613 addition in all of the cell lines tested (Fig. 7A). This change in membrane potential was not caused by a general increase in membrane permeability. Breast cancer cell lines were more reactive than HMLE cells (Fig. 7B). Low levels of induction of depolarisation by ZG613 in HMLE cells could be related to a rather positive basal resting membrane potential value comparing to human mammary epithelia (HMEC)^74^. Regarding HMLETwist cell line, enhanced sensitivity to plasma membrane permeabilisation may level the net changes in membrane potential. Mechanistic study of ion fluxes, MIFE, revealed that ZG613 barely affects Na^+^ and K^+^ transport through the plasma membrane in breast cancer and EMT cells (Fig. 8A, Fig S2), which confirmed MD simulations findings. We previously concluded that adamantane side arm in ZG613 enhances conformational flexibility and increases affinity for binding of monovalent cations^40^. On the other hand, strong binding may prevent release of the ions from the crowns^67^, thus it may interfere with ion release from ZG613 once the membrane is crossed. However, according to the MD simulation in POPC membrane, K^+^ dissociates from ZG613 and returns to the bulk water (video 3, Supporting information), which could indicate that the presence of the hydrophobic adamantane groups is not sufficient to overcome the energetic penalty to drag a charged species deep inside the bilayer. Another possible explanation of the ion transport failure by ZG613, according to the MD simulation, may lie in the formation of aggregates on the water side that interfere with ion transport, but still causes physical disruption of the plasma membrane. However, since ZG613 depolarises cells, we reasoned that the compound may cause an increase in intracellular Ca^2+^ levels. Increase in the intracellular calcium is regarded a key event after membrane damage^1^. We caught moderate but consistent Ca^2+^ rise as soon as four minutes after the addition of ZG613, and furthermore, Ca^2+^ levels increased even when cells were placed in medium without Ca^2+^ (Fig. 8B). This could indicate that ZG613 depletes ER Ca^2+^ stores, possibly by disruption of ER membranes.

To clarify whether and at what concentration ZG613 is toxic to mice, and furthermore, whether ZG613 is able to inhibit tumour growth *in vivo*, we had to overcome the compound’s high hydrophobicity, a major difficulty for obtaining high concentrations in nontoxic solvents. We chose Corn oil, since it is successfully used as a solvent for i.p. administration of hydrophobic drugs such as Tamoxifen in rodents^83^. The highest obtained dose was still much lower the predicted LD_50_ dose. Nonetheless, we observed a mild retardation of tumour growth and decrease in formations of metastasis (Fig. 9). In general, the effects *in vivo* were milder than anticipated, most likely due to the low solubility of the compound, which represents significant limitation for the potential therapeutic applications.

## Conclusions

Our proprietary adamantane derived diaza-crown ether ZG613 targets breast cancer cells and breast EMT-model cells by physical disruption of plasma membrane and impairments of mitochondrial functions rather than by disturbance in ion transport across the plasma membrane. Since EMT-model cells exhibited sensitivity to ZG613, this model represents a useful tool to study drugs that induce changes in plasma membrane integrity and mitochondrial alterations.

Thus, we proposed membrane integrity and mitochondrial functions as fitting targets in EMT, and potentially CSCs, among tumours, using small hydrophobic compounds as adamantyl crown ether ZG613.

## Acknowledgments

This study was supported by Croatian Science Foundation project (number IP-2013-5660 “MultiCaST”) to MK. The work was also supported by the FP7-REGPOT-2012-2013-1 project (grant agreement number 316289 – InnoMol) and by the COST Action BM1406.

## Author contributions

Katja Ester and Marijeta Kralj conceived and initiated this project. Katja Ester was responsible for conceptualization, methodology, data analysis, experimental investigation and writing original draft manuscript. Marija Mioč, Pavel Spurny, Daniel Bonhenry, Marko Marjanović, Lidija Uzelac, Jelena Gabrilo, Wolfgang Schreibmayer, Babak Minofar and Jost Ludwig contributed to the methodology, experimental investigation, data analysis or manuscript revision. Kata Majerski designed and Tatjana Šumanovac synthesized the compound. Marijeta Kralj made contributions in the conceptualization, manuscript revision and funding support.

