## Supplementary Information for "Targeting plasma membrane and mitochondrial instability in breast cancer cells and breast epithelial to mesenchymal transition-model cells by adamantyl diaza-crown ether ZG613"

**Table S1**. Significantly up-regulated and down-regulated genes among differentially expressed gens in HMLEpBp and HMLETwist cells induced by ZG613

| HMLEpBp |  |  |  |
| --- | --- | --- | --- |
| Gene | log2(Fold Change) | Std. Err. log2(Fold Change) | Significant |
| CXCL8 | 0,74 | 0,17 | TRUE |
| ACAT2 | 0,65 | 0,13 | TRUE |
| IGFBP3 | 0,63 | 0,15 | TRUE |
| VGLL3 | 0,59 | 0,16 | TRUE |
| SERPINB2 | 0,57 | 0,15 | TRUE |
| AKAP12 | 0,54 | 0,13 | TRUE |
| RAB3B | 0,54 | 0,14 | TRUE |
| ABL2 | 0,49 | 0,13 | TRUE |
| ALDH1A3 | 0,48 | 0,13 | TRUE |
| F3 | 0,42 | 0,11 | TRUE |
| VPS51 | -0,5 | 0,14 | TRUE |
| ARHGEF16 | -0,51 | 0,14 | TRUE |
| HSPBP1 | -0,51 | 0,14 | TRUE |
| SAPCD2 | -0,52 | 0,14 | TRUE |
| C7orf50 | -0,52 | 0,15 | TRUE |
| PIM3 | -0,53 | 0,14 | TRUE |
| CEBPB | -0,53 | 0,14 | TRUE |
| TNNI2 | -0,53 | 0,14 | TRUE |
| MRPL57 | -0,53 | 0,15 | TRUE |
| ZNF598 | -0,53 | 0,15 | TRUE |
| DDIT4 | -0,54 | 0,12 | TRUE |
| MXD4 | -0,54 | 0,14 | TRUE |
| NGFR | -0,54 | 0,15 | TRUE |
| NOSIP | -0,54 | 0,15 | TRUE |
| GSDMD | -0,54 | 0,15 | TRUE |
| AQP3 | -0,55 | 0,14 | TRUE |
| GPX1 | -0,55 | 0,14 | TRUE |
| TMEM129 | -0,55 | 0,15 | TRUE |
| CCDC124 | -0,55 | 0,15 | TRUE |
| H1FX | -0,55 | 0,15 | TRUE |
| RHOT2 | -0,56 | 0,16 | TRUE |
| PGAP3 | -0,56 | 0,16 | TRUE |
| UNC5B | -0,57 | 0,14 | TRUE |
| GTPBP6 | -0,57 | 0,16 | TRUE |
| SIVA1 | -0,57 | 0,16 | TRUE |
| IGFBP2 | -0,57 | 0,16 | TRUE |
| POLD1 | -0,58 | 0,15 | TRUE |
| ATP6V0C | -0,58 | 0,15 | TRUE |
| ATP5D | -0,58 | 0,16 | TRUE |
| MICALL2 | -0,58 | 0,16 | TRUE |
| COL18A1 | -0,59 | 0,14 | TRUE |
| CBX4 | -0,59 | 0,15 | TRUE |
| MRPL34 | -0,59 | 0,15 | TRUE |
| WDR18 | -0,59 | 0,15 | TRUE |
| CDKN2A | -0,59 | 0,16 | TRUE |
| TELO2 | -0,59 | 0,16 | TRUE |
| NPDC1 | -0,59 | 0,17 | TRUE |
| ZNF205 | -0,59 | 0,17 | TRUE |
| FLYWCH2 | -0,59 | 0,17 | TRUE |
| LSM7 | -0,6 | 0,15 | TRUE |
| MBD3 | -0,6 | 0,15 | TRUE |
| CHTF18 | -0,6 | 0,16 | TRUE |
| GPC1 | -0,6 | 0,16 | TRUE |
| SSBP4 | -0,6 | 0,17 | TRUE |
| ARVCF | -0,6 | 0,17 | TRUE |
| WNT10A | -0,61 | 0,15 | TRUE |
| RECQL4 | -0,61 | 0,15 | TRUE |
| ZDHHC24 | -0,61 | 0,17 | TRUE |
| NUBP2 | -0,62 | 0,15 | TRUE |
| CROCC | -0,62 | 0,16 | TRUE |
| NDUFS7 | -0,62 | 0,16 | TRUE |
| PITX1 | -0,62 | 0,16 | TRUE |
| LRRC45 | -0,62 | 0,16 | TRUE |
| ULK1 | -0,63 | 0,14 | TRUE |
| ZNF219 | -0,63 | 0,16 | TRUE |
| MEX3D | -0,63 | 0,16 | TRUE |
| CHAC1 | -0,63 | 0,18 | TRUE |
| TPGS1 | -0,63 | 0,18 | TRUE |
| ABCA7 | -0,64 | 0,16 | TRUE |
| C9orf142 | -0,64 | 0,16 | TRUE |
| PIDD1 | -0,64 | 0,16 | TRUE |
| CEBPD | -0,64 | 0,18 | TRUE |
| PKD1P1 | -0,65 | 0,14 | TRUE |
| ALDH16A1 | -0,65 | 0,16 | TRUE |
| PWWP2B | -0,65 | 0,17 | TRUE |
| TRAPPC5 | -0,65 | 0,17 | TRUE |
| C19orf60 | -0,65 | 0,18 | TRUE |
| ANKRD9 | -0,65 | 0,18 | TRUE |
| TSPO | -0,67 | 0,16 | TRUE |
| MESDC1 | -0,67 | 0,16 | TRUE |
| C11orf31 | -0,67 | 0,17 | TRUE |
| C16orf93 | -0,67 | 0,18 | TRUE |
| MACROD1 | -0,67 | 0,19 | TRUE |
| FGFR3 | -0,68 | 0,13 | TRUE |
| E4F1 | -0,68 | 0,16 | TRUE |
| SEMA6B | -0,68 | 0,18 | TRUE |
| SYTL1 | -0,69 | 0,15 | TRUE |
| MFSD3 | -0,69 | 0,16 | TRUE |
| IRX4 | -0,69 | 0,16 | TRUE |
| FAM195A | -0,69 | 0,17 | TRUE |
| PPP1R16A | -0,7 | 0,17 | TRUE |
| JUND | -0,7 | 0,17 | TRUE |
| PHLDA2 | -0,7 | 0,17 | TRUE |
| RHPN1 | -0,7 | 0,18 | TRUE |
| ABHD17A | -0,71 | 0,16 | TRUE |
| GDF15 | -0,71 | 0,18 | TRUE |
| VASN | -0,73 | 0,17 | TRUE |
| SCAND1 | -0,73 | 0,17 | TRUE |
| KREMEN2 | -0,73 | 0,17 | TRUE |
| JOSD2 | -0,73 | 0,18 | TRUE |
| PLEKHH3 | -0,74 | 0,16 | TRUE |
| FAM173A | -0,74 | 0,18 | TRUE |
| PDF | -0,74 | 0,18 | TRUE |
| MXD3 | -0,75 | 0,18 | TRUE |
| PRR7 | -0,75 | 0,18 | TRUE |
| NME2 | -0,75 | 0,19 | TRUE |
| MIB2 | -0,77 | 0,17 | TRUE |
| HAGHL | -0,77 | 0,18 | TRUE |
| NME3 | -0,77 | 0,18 | TRUE |
| SYT8 | -0,78 | 0,14 | TRUE |
| JUNB | -0,79 | 0,17 | TRUE |
| RTN4R | -0,79 | 0,17 | TRUE |
| MZT2A | -0,8 | 0,17 | TRUE |
| ID1 | -0,81 | 0,14 | TRUE |
| C4orf48 | -0,81 | 0,18 | TRUE |
| MZT2B | -0,85 | 0,17 | TRUE |
| SNHG19 | -0,86 | 0,17 | TRUE |
| MIF | -0,9 | 0,17 | TRUE |
| COMTD1 | -0,91 | 0,17 | TRUE |
| FBXL15 | -0,93 | 0,18 | TRUE |
| CCDC85B | -1,14 | 0,19 | TRUE |

| HMLETwist |  |  |  |
| --- | --- | --- | --- |
| Gene | log2(Fold Change) | Std. Err. log2(Fold Change) | Significant |
| MVD | 1,02 | 0,16 | TRUE |
| CAPN15 | 0,96 | 0,17 | TRUE |
| SCAND1 | 0,9 | 0,19 | TRUE |
| LRFN4 | 0,89 | 0,18 | TRUE |
| FABP3 | 0,86 | 0,18 | TRUE |
| ABHD17A | 0,85 | 0,19 | TRUE |
| C19orf60 | 0,85 | 0,2 | TRUE |
| C4orf48 | 0,84 | 0,18 | TRUE |
| ZNF628 | 0,84 | 0,2 | TRUE |
| TPGS1 | 0,83 | 0,19 | TRUE |
| PRKCDBP | 0,83 | 0,19 | TRUE |
| PRR7 | 0,83 | 0,19 | TRUE |
| TRAPPC5 | 0,81 | 0,19 | TRUE |
| PHLDA2 | 0,8 | 0,19 | TRUE |
| JUND | 0,79 | 0,19 | TRUE |
| PPP1R16A | 0,78 | 0,18 | TRUE |
| FAM173A | 0,78 | 0,2 | TRUE |
| ZNF865 | 0,77 | 0,2 | TRUE |
| RNF126 | 0,76 | 0,17 | TRUE |
| CHTF18 | 0,76 | 0,18 | TRUE |
| BCAR1 | 0,75 | 0,16 | TRUE |
| TWIST1 | 0,75 | 0,16 | TRUE |
| MZT2A | 0,75 | 0,19 | TRUE |
| KLHL17 | 0,75 | 0,19 | TRUE |
| CROCC | 0,75 | 0,19 | TRUE |
| PLEC | 0,74 | 0,15 | TRUE |
| TLX2 | 0,74 | 0,2 | TRUE |
| ZDHHC8 | 0,73 | 0,17 | TRUE |
| VASN | 0,73 | 0,18 | TRUE |
| H1FX | 0,73 | 0,18 | TRUE |
| NME2 | 0,73 | 0,2 | TRUE |
| C20orf24 | 0,73 | 0,2 | TRUE |
| ATP6V0C | 0,72 | 0,17 | TRUE |
| SBNO2 | 0,72 | 0,17 | TRUE |
| ATP5D | 0,72 | 0,18 | TRUE |
| TM7SF2 | 0,72 | 0,19 | TRUE |
| PABPN1 | 0,72 | 0,19 | TRUE |
| CLEC11A | 0,72 | 0,2 | TRUE |
| ZNF787 | 0,7 | 0,18 | TRUE |
| H2AFX | 0,7 | 0,18 | TRUE |
| DHRS4 | 0,7 | 0,19 | TRUE |
| MAP1S | 0,69 | 0,18 | TRUE |
| FASN | 0,68 | 0,14 | TRUE |
| DOT1L | 0,68 | 0,16 | TRUE |
| SCAF1 | 0,68 | 0,16 | TRUE |
| MMP17 | 0,68 | 0,18 | TRUE |
| SDF2L1 | 0,68 | 0,18 | TRUE |
| MRPS24 | 0,68 | 0,18 | TRUE |
| NCOR2 | 0,67 | 0,16 | TRUE |
| EPN1 | 0,67 | 0,17 | TRUE |
| SSBP4 | 0,67 | 0,18 | TRUE |
| SCRIB | 0,66 | 0,17 | TRUE |
| PRR12 | 0,66 | 0,17 | TRUE |
| RFX1 | 0,66 | 0,18 | TRUE |
| CHST12 | 0,66 | 0,18 | TRUE |
| SLC2A6 | 0,65 | 0,16 | TRUE |
| FAM83H | 0,65 | 0,17 | TRUE |
| PKMYT1 | 0,64 | 0,15 | TRUE |
| TERT | 0,64 | 0,16 | TRUE |
| TMEM259 | 0,64 | 0,17 | TRUE |
| TRABD | 0,64 | 0,18 | TRUE |
| RPUSD1 | 0,63 | 0,16 | TRUE |
| MFSD10 | 0,63 | 0,17 | TRUE |
| UBE2S | 0,62 | 0,16 | TRUE |
| JUN | 0,61 | 0,16 | TRUE |
| LENG8 | 0,6 | 0,16 | TRUE |
| TGFB1 | 0,59 | 0,16 | TRUE |
| REPIN1 | 0,59 | 0,16 | TRUE |
| SLC52A2 | 0,59 | 0,16 | TRUE |
| MFSD12 | 0,58 | 0,15 | TRUE |
| DHCR7 | 0,58 | 0,16 | TRUE |
| ID1 | 0,57 | 0,15 | TRUE |
| CLCN7 | 0,56 | 0,15 | TRUE |
| PIEZO1 | 0,54 | 0,15 | TRUE |
| ARHGDIA | 0,52 | 0,14 | TRUE |
| KRT6A | -0,62 | 0,17 | TRUE |

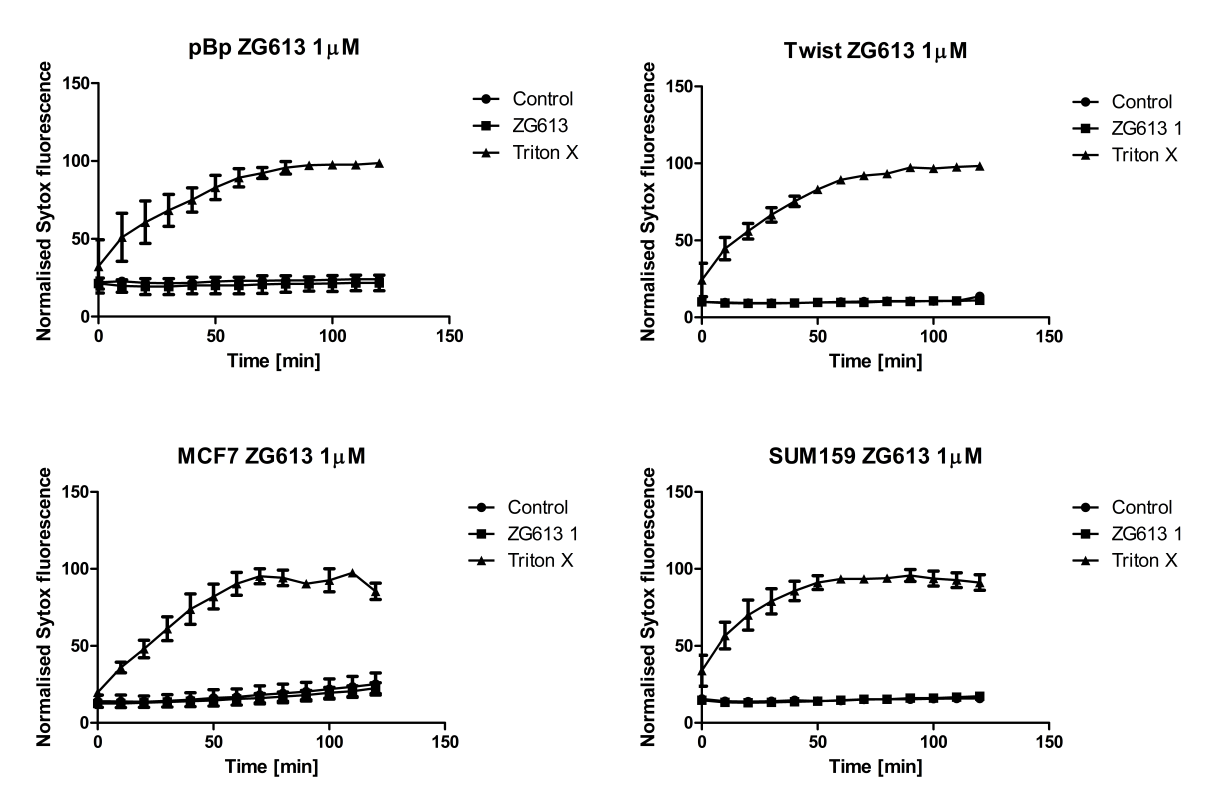

**Figure S1** Time course of Sytox influx induced by ZG613 is presented. Permeation of Sytox dye was monitored in breast cancer and breast EMT-model cell lines up to 120 min after the addition of 1 µM ZG613, DMSO or Triton X. Fluorescence was measured every 10 min using Tecan M200 microplate reader. Plots present data from at least three independent experiments. Maximal value obtained with non-ionic detergent Triton X (0.3 %) was set as 100%.

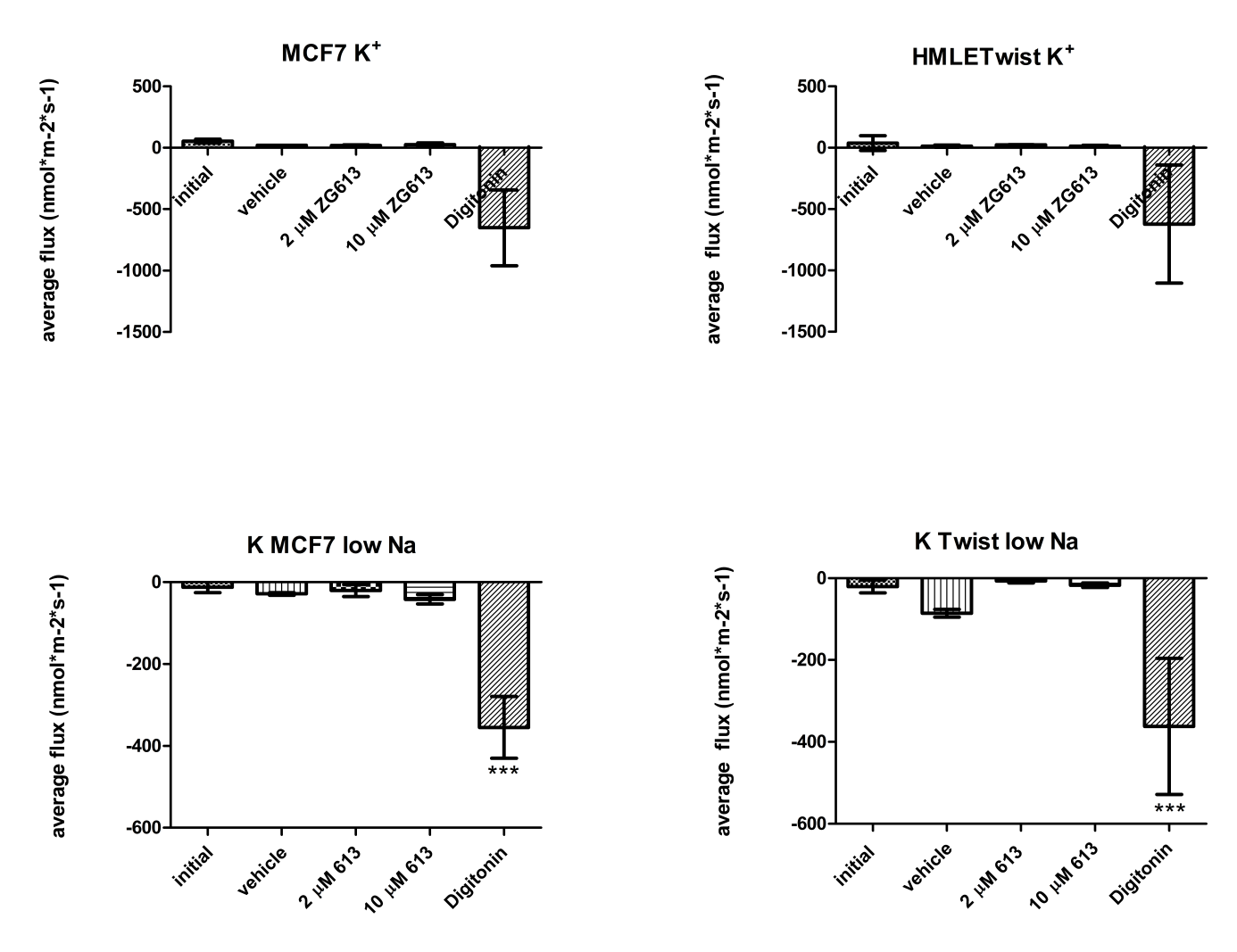

**Figure S2** Histograms represent average fluxes of K^+^ induced by ZG613, vehicle (DMSO) and non-ionic detergent Digitonin in HMLETwist and MCF7 cell lines. High Na^+^: measuring solution with physiological Na+ concentration (137mM). Low Na^+^: measuring solution with low Na+ concentration (0.2 mM). Data were analysed using 1 way ANOVA with Tukey’s Multiple comparison test; *** p<0.005
